# Loss of Diurnal Oscillatory Rhythms in Gut Microbiota Correlates with Progression of Atherosclerosis

**DOI:** 10.1101/2024.09.23.614533

**Authors:** He Zhang, Xiaohan Zhang, Zihan Yun, Yang Chen, Suhua Cang, Yating Shao, Erteng Jia, Renjin Chen

## Abstract

Circadian rhythms in gut microbiota composition are crucial for metabolic function and disease progression, yet the diurnal oscillation patterns of gut microbiota in atherosclerotic cardiovascular disease (ASCVD) and their role in disease progression remain unknown. Here, we investigate gut bacterial dynamics in ApoE^-/-^ mice within a day, and elucidated the dynamic changes in fecal microbiota composition and function differences among C57BL/6 and ApoE^-/-^ mice with standard chow diet or high-fat, high-cholesterol diet under ad libitum conditions. Compared with C57BL/6 mice, ApoE^-/-^ mice exhibit significant differences in fecal microbial composition. Rhythmic analysis showed that the dynamic changes in the composition and function of fecal microbiota in ApoE^-/-^ mice were significantly different from those in C57BL/6 mice. We further found that the rhythmic strains (*Blautia Coccoides*) inhibit the progression of ASCVD by improving the intestinal and endothelial barrier function. Our findings demonstrate that diurnal oscillations in gut microbiota are closely related to the progression of ASCVD, and provide a new insight for microbial-targeted therapies for ASCVD.

## Introduction

Cardiovascular diseases (CVDs) are the most common non-communicable diseases worldwide, accounting for about one-third of global deaths ^[1]^. CVDs are the main cause of death among Asians, have become a public health problem that seriously threatens the health of people, and have brought huge economic burdens to Asian countries ^[2]^. Atherosclerotic cardiovascular disease (ASCVD), whose clinical symptoms are lipid accumulation and inflammation of the aorta, is the main cause of CVDs ^[3]^. In recent years, an unhealthy diet has become the most significant factor in the occurrence and death of CVDs in Chinese ^[4]^. However, previous studies on the regulation of dietary behavior in the progression of CVDs have mainly relied on epidemiological investigations, failing to conduct in-depth research and elucidate its underlying mechanisms.

Among the 15 risk factors significantly associated with CVDs, an unhealthy diet is the second major risk factor, after hypertension ^[5]^. Dietary behavior is closely related to the composition and function of gut microbiota. Interactions between gut microbiota composition, diet, and host can influence the progression of ASCVD. Metabolites of gut microbiota, such as trimethylamine oxide ^[6]^, short-chain fatty acids ^[7]^, secondary bile acids ^[8]^, indole-3-propionic acid ^[9]^, and phenylacetylglutamine ^[10]^, significantly affect the progression of ASCVD, indicating that metabolites derived from gut microbiota can regulate health and disease processes individually or collectively through host metabolic pathways.

The gut microbiome is closely related to circadian rhythms, and dysregulation of circadian genes as well as feeding time are known disruptors of the gut microbiome in animal models ^[11, 12]^. Environmental factors, especially dietary composition, timing of intake, antibiotic usage, and the administration of other medications, significantly drive the dynamics of gut microbiota^[13]^. Dietary composition and patterns play a crucial role in shaping the composition and function of gut microbiota and are the easiest factors to implement interventions ^[14]^. Under normal dietary conditions, analyzing the composition and functional changes of gut microbiota in patients is of great significance for using personalized nutritional interventions to reshape the interaction between host and microbiota, thereby controlling and preventing disease progression, and contributing to the development of novel therapeutic strategies targeting the microbiota of ASCVD. To test the diurnal oscillatory pattern of microbiota in ApoE^-/-^ mice, the diurnal fecal microbiota patterns were examined in mice fed ad libitum with a chow diet or a high-fat, high-cholesterol diet (HFHCD).

## Materials and methods

### Experimental animals, design, and diet

C57BL/6 mice and ApoE^-/-^ mice on a C57BL/6 background were purchased from Jiangsu GemPharmatech Co., Ltd. All animal experiments were approved by the Animal Care and Utilization Committee of Xuzhou Medical University. For the ad libitum feeding experiments, six 6-week-old C57BL/6 or ApoE^-/-^ mice (three females and three males) were randomly assigned to either the standard chow diet group or the HFHCD group, with three female mice and three male mice in each group. After 12-week administration with a strict 12 h/12 h light/dark cycle after shipment (lights are on from ZT8 to ZT20), we obtained fecal samples every 6 h within a 24-h period at four different time points of 05:00 (ZT5), 11:00 (ZT11), 17:00 (ZT17) or 23:00 (ZT23). For the microbial interference experiment, eighteen 5-week-old ApoE^-/-^ male mice were randomly divided into three groups: the PBS group, the Ina-B.Coccides group, and the B.Coccides group. After two weeks of interference with broad-spectrum antibiotics (ampicillin, neomycin, metronidazole, and vancomycin), mice in the B.Coccides group received 0.2 mL of bacterial solution (5x10^9 CFU) three times each week, the Ina-B.Coccides group received 0.2 mL of autoclaved bacterial solution, and the PBS group received an equal volume of PBS. During the experiments, mice had free access to a high-fat, high-cholesterol diet and water. Oral gavage administration began, and the body weight and feed intake of the mice were monitored daily. After eleven weeks, the colon contents, submandibular vein blood, and aortic tissue were collected. In addition, subcutaneous inguinal adipose tissue (IAT) and epididymal adipose tissue (EAT) were collected and weighed.

### Cell culture

Immortalized mouse aortic endothelial cells (MAEC; FH-042YSH) were purchased from Shanghai Fuheng Biotechnology Co., Ltd. MAEC cells were cultured in mouse aortic endothelial cell complete medium (PYM018; Shanghai Fuheng Biotechnology Co., Ltd.), which contained 1% (v/v) penicillin/streptomycin (100 Units/ml penicillin and 100 µg/ml streptomycin).

Gallic acid (APExBIO; USA) was dissolved to 1 µg/µL in sterile water. To evaluate the effects of gallic acid on aortic barrier function and cholesterol transport in mice, the culture medium was replaced with 2 mL of fresh complete medium + saline (6 wells) or complete medium containing GA (2 µL, 10 µL, or 50 µL). The plates were then incubated for 24 hours. At the end of the culture period, samples were collected for gene quantification and Western blot analysis.

### Full-length 16S sequencing

Microbial DNA was isolated from mouse fecal samples using the E.Z.N.A.® stool DNA Kit (Omega Bio-tek, Norcross, GA, U.S.) following the manufacturer’s protocols. The V1-V9 region of the bacterial 16S ribosomal RNA gene was amplified by PCR (95 °C for 2 min, then 27 cycles at 95 °C for 30 s, 55 °C for 30 s, and 72 °C for 60 s, with a final extension at 72 °C for 5 min) using primers 27F 5’-AGRGTTYGATYMTGGCTCAG-3’ and 1492R 5’-RGYTACCTTGTTACGACTT-3’, with a unique eight-base barcode for each sample.

### 16S sequencing data processing and analysis

SMRTbell libraries were prepared from the amplified DNA through blunt-ligation following the manufacturer’s protocols. (Pacific Biosciences). Purified SMRTbell libraries from both the Zymo and HMP mock communities were sequenced on dedicated PacBio Sequel II 8M cells utilizing the Sequencing Kit 2.0 chemistry. The purified SMRTbell libraries from the pooled and barcoded samples were sequenced on a single PacBio Sequel II cell. Shanghai Biozeron Biotechnology Co. Ltd (Shanghai, China) performed all amplicon sequencing. SMRT Link Analysis software (version 9.0) was used to process PacBio raw reads, and SMRT Portal was used to filter low-quality score bases. If they contained 10 consecutive identical bases, the barcode, primer sequences, and chirmas will be removed. OTUs were clustered with UPARSE (version 7.1, http://drive5.com/uparse/) using a similarity threshold of 98.65%, and chimeric sequences were identified and removed using UCHIME. The RDP Classifier (http://rdp.cme.msu.edu/) was employed to classify the phylogenetic affiliation of each 16S rRNA gene sequence, with analysis conducted in the Silva (SSU132) 16S rRNA database and a confidence threshold set at 70% ^[15]^. The diversity indices, including the Observe, Chao1, ACE, Shannon, Simpson, and Pielou were conducted by the R package (MicrobiotaProcess, version 1.13.2.994). Beta diversity analysis was conducted using UniFrac ^[16]^ to compare the results of principal component analysis (PCA) from the community ecology package R-forge, which generated PCA graphs with the Vegan 2.0 package. Analysis and visualization of microbial community structure was conducted using by R package (MicrobiotaProcess, version 1.13.2.994). Linear discriminant analysis Effect Size (LEfSe) analysis was conducted using by R package (microbiomeMarker, version 1.6.0). PICRUSt2 Analysis was performed using the OmicStudio Analysis at https://www.omicstudio.cn/analysis/.

### Culture and administration of Blautia coccoides

*Blautia coccoides* (ATCC 29236, American Type Culture Collection, Manassas, VA) were cultured anaerobically in modified chopped meat medium, ATCC medium 1490. The concentration of bacteria was determined by measuring the absorbance at a wavelength of 600 nm. Then, 5×10^9^ cfu of *Blautia coccoides* in 0.2 mL PBS was orally gavaged three times each week to ApoE^−/−^ mice. In the Ina-B.Coccides group, *Blautia coccoides* were heat killed at 121°C under 225-kPa pressure for 15 minutes.

## LC-MS/MS

The *Blautia coccoides* culture supernatant (100 μL) or bacteria-cultured medium (100 μL) was thoroughly mixed with pre-cooled methanol (400 μL) and vortexed vigorously. After incubation on ice for 5 min, the sample was centrifuged at 15,000 rpm and 4°C for 5 min. Then, the supernatant was diluted with LC-MS grade water to a final concentration of 53% methanol. Next, the sample was transferred to a new Eppendorf tube and centrifuged at 15000 g and 4°C for 10 min. Finally, the supernatant was injected into the LC-MS/MS system for analysis. UHPLC-MS/MS analysis was conducted at Biozeron Co., Ltd. (Shanghai, China) using a Vanquish UHPLC system (Thermo Fisher, Germany) and the Orbitrap Q ExactiveTM HF mass spectrometer (Thermo Fisher, Germany), as previously described ^[17]^.

### Atherosclerotic plaque analysis

To evaluate the lipid deposition in whole arteries, the entire “aortic tree” fixed in 10% neutral buffered formalin was stained withSudan IV. Aortic images were captured with Leica EZ 4W (Leica, Germany). The aortic root was embedded into an OCT embedding agent, sliced into frozen sections (approximately 5-µm thick), and stored at −20°C. For analysis of the lesion area, Oil red O staining and Hematoxylin and eosin (H&E) staining for frozen sections were performed. The quantification of lesion area and size was quantitatively calculated using image analysis software (Image-Pro Plus-6, Media Cybernetics, Inc).

### Biochemical parameters of plasma

Triglyceride (TG), total cholesterol (T-CHO), high-density lipoprotein-cholesterol (HDL-C), and low-density lipoprotein-cholesterol (LDL-C) concentrations in plasma of ApoE^-/-^ mice were determined using an Olympus AU400 Automatic Biochemical Analyzer (Tokyo, Japan).

### Real-time quantitative PCR analysis

Total RNA was extracted from MAEC under the guidance of the manufacturer of the RNeasy mini kit (Qiagen, Hilden, North Rhine-Westphalia, Germany). Reverse transcribed 1 μg RNA to cDNA using PrimeScript™ RT Master Mix kit (TaKaRa Co. Ltd. Dalian, China). qPCR was performed on the relative expression levels of target genes by ABI 7300 sequence detector (SDS, Foster City, CA, USA) using the SYBR Green PCR kit (TaKaRa, Co., Dalian, China). Invitrogen Life Technologies (Invitrogen, Shanghai, China) manufactured all of the primers, and their sequences are provided in Table A2. The formula 2^-ΔΔCt^ was used to compute the relative quantification of gene expression differences ^[18]^.

### Western blot analysis

Total proteins were extracted from the mouse colon and MAEC using RIPA lysis buffer (Thermo Scientific, Waltham, MA, USA) including protease inhibitors and phosphatase inhibitors. Protein concentrations were determined using the BCA protein assay kit (Nanjing Jiancheng Institute of Bioengineering, Nanjing), 20 μg of isolated proteins were isolated by 10% SDS PAGE gel electrophoresis, and then transferred to a polyvinylidene fluoride (PVDF) membrane (Millipore, Bedford, MA, USA). Antibodies against: β-Actin (bs-0061R, 1: 1000, Bioss), NF-κB p65 antibody (bsm-33117M, 1: 800, Bioss), Occludin antibody (bs-10011R, 1: 500, Bioss), ZO-1 antibody (bs-34023R, 1: 500, Bioss), Claudin 5 antibody (bs-10296R, 1: 500, Bioss), CLDN1 antibody (bs-1428R1: 500, Bioss), CD144 (VE-cadherin) antibody (14-1441-82, 1: 500, Invitrogen).

### Statistical analysis

Statistical analyses and chart drawing were performed using R (version 4.4.1), SPSS 26.0 (SPSS Inc., Chicago, IL, USA), and GraphPad Prism 9.5 (GraphPad Software, San Diego, CA, USA). The R package JTK_circle analysis method was employed to detect diurnal fluctuations with a 24-hour cycle and 6-hour intervals ^[19, 20]^. Differences in microbial abundance (at the phylum and genus levels) were analyzed using the Kruskal-Wallis test. Function of the gut microbiota between the light phase (ZT11, and ZT17) and the dark phase (ZT5 and ZT23) was analyzed using the Mann-Whitney U-test. Diversity index, abundance index, gene and protein expression, and plasma biochemical parameters were detected by one-way ANOVA analysis, followed by Duncan’s test.

## Results

### Significant differences in the fecal microbiota composition between ApoE^-/-^ mice and C57BL/6 mice at different time points within a day

To investigate the structural changes in the gut microbiota associated with AS, we collected feces samples from C57_Chow, C57_HFD, ApoE^-/-^_Chow, and ApoE^-/-^_HFD mice after 12-week period under an *ad libitum* feeding regimen, and performed full-length 16S rRNA sequencing. There were significant differences in the diversity index and composition of fecal microbiota among the four groups of mice (Fig. S1). To reveal the structural differences in the fecal microbiota between ApoE^-/-^ and C57BL/6 mice, we further analyzed the fecal microbiota composition among the four groups of mice at different time points within the day. In the alpha diversity analysis, compared to the C57_Chow group, the ApoE^-/-^_HFD group did display significant differences in alpha diversity at ZT5, ZT11, and ZT23, including Chao1 and ACE (Fig 1). In comparison to the C57_HFD group, the ApoE^-/-^_Chow and ApoE^-/-^_HFD groups did display significant differences in alpha diversity, including Chao1, ACE, Shannon, and Simpson (Fig 1). The results of PCoA showed that there were significant differences in the composition of fecal microbiota in the four groups at different time points within a day, indicating that the fecal microbial composition of these four groups of mice could be significantly distinguished regardless of the time point of observation.

**Figure 1.**
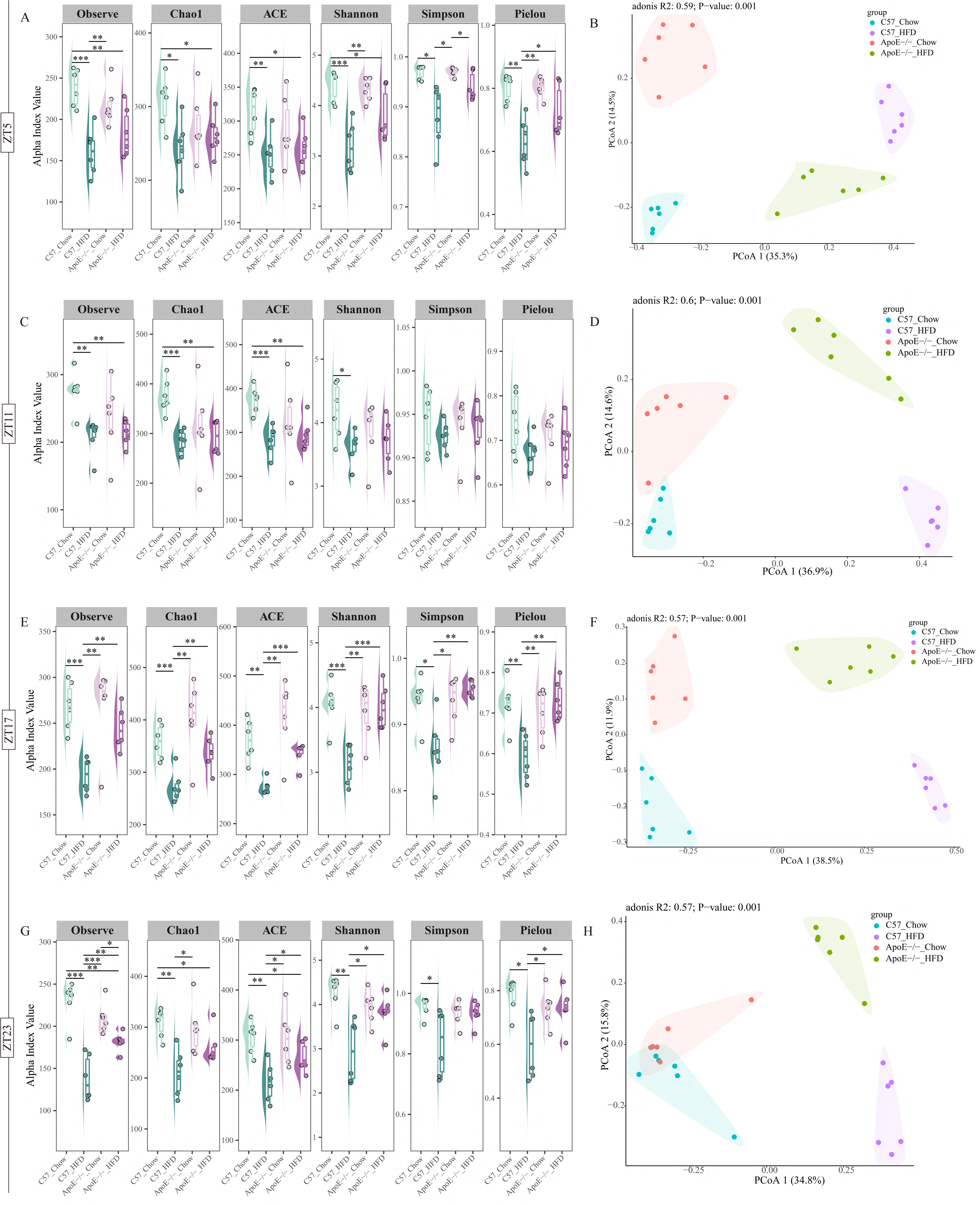
α-diversity and β-diversity of the fecal microbiota of C57_Chow, C57_HFD, ApoE-/-_Chow, and ApoE-/-_HFD mice at different time points. (A, C, E, G) α-diversity of fecal microbiota at ZT5, ZT11, ZT17, and ZT23; (B, D, F, H) PCoA of fecal microbiota at ZT5, ZT11, ZT17, and ZT23. Relative variable importance (R2) and significance (*P*) were calculated by PERMANOVA (Adonis) analysis. ZT, zeitgeber; * indicates *P* < 0.05, ** indicates *P* < 0.01, *** indicates *P* < 0.001; n=6.

To compare the differences in fecal microbiota composition among four groups of mice, we employed non-parametric tests for analyzing the relative abundance of fecal microbiota. The results revealed distinct microbial taxa associated with each group. The relative abundance of fecal microbiota in the four groups of mice exhibited significant variations at both the phylum and genus levels at different time points within a day (Fig 5A-5D, 5H-5K). These findings suggested significant differences in fecal microbiota composition between ApoE^-/-^ mice and C57BL/6 mice at different time points within a day, and that fecal microbiota composition exhibited dynamic changes (Fig 2).

**Figure 2.**
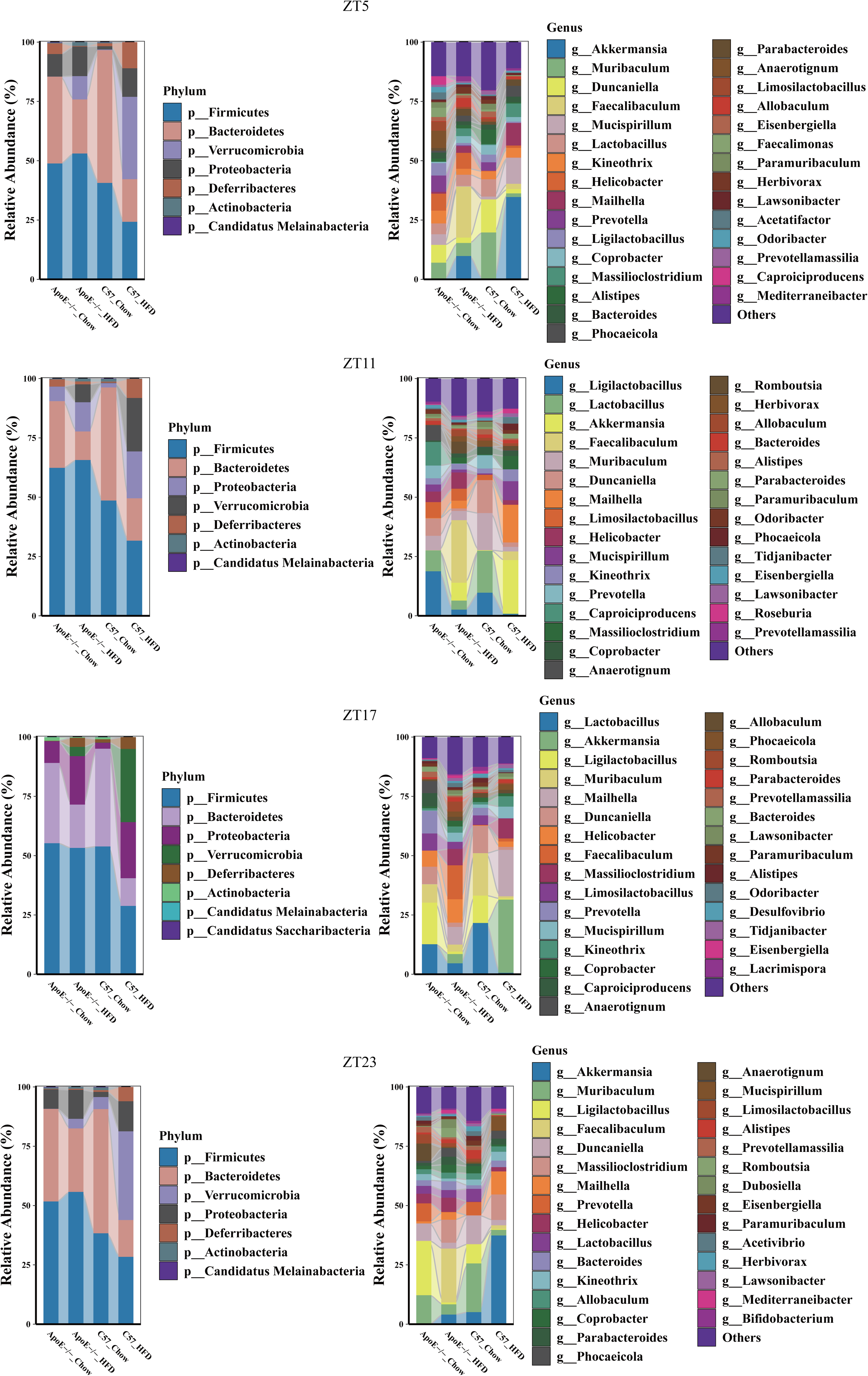
Composition of the fecal microbial taxa at the phylum and genus level of C57_Chow, C57_HFD, ApoE-/-_Chow, and ApoE-/-_HFD mice at different time points. (A, C, E, G) Barplot of fecal microbiota at the phylum at ZT5, ZT11, ZT17, and ZT23. (B, D, F, H) Barplot of fecal microbiota at the genus at ZT5, ZT11, ZT17, and ZT23. ZT, zeitgeber; * indicates *P* < 0.05, ** indicates *P* < 0.01, *** indicates *P* < 0.001; n=6.

### Diurnal fluctuation of fecal microbiota composition of ApoE^-/-^ mice and C57BL/6 mice

To elucidate the dynamic changes in the structure of fecal microbiota in ApoE^-/-^ mice, we conducted a comprehensive analysis of the changes in the composition of fecal microbiota in four groups of mice at different time points within a day. PCoA analysis based on Bray-Curtis dissimilarity differentiated fecal samples from ApoE^-/-^ group mice at different time points overall (R2 = 0.19, *P*= 0.029, Fig 3F). Further analysis was conducted on the fecal microbiota composition in four groups of mice at different time points (Fig 4). At the phylum level, C57_Chow mice exhibited a higher abundance of Bacteroidetes at ZT5 and ZT23, and a higher abundance of Firmicutes at ZT11 and ZT17. C57_HFD mice exhibited a higher abundance of Verrucomicrobia at ZT5, ZT17, and ZT23, and a higher abundance of Firmicutes at ZT11. ApoE^-/-^_Chow and ApoE^-/-^_HFD mice exhibited a higher abundance of Firmicutes at different time points within a day. Similarly, at the genus level, C57_Chow mice exhibited a higher abundance of *Muribaculum* at ZT5 and ZT23, and a higher abundance of *Lactobacillus* at ZT11 and ZT17. C57_HFD mice exhibited a higher abundance of *Akkermansia* at different time points within a day. ApoE^-/-^_Chow mice exhibited a higher abundance of *Duncaniella* at ZT5, and a higher abundance of *Ligilactobacillus* at ZT11, ZT17, and ZT23. C57_HFD mice exhibited a higher abundance of *Faecalibaculum* at different time points within a day. These data suggested that the fecal microbial composition exhibited rhythmic oscillations within a day.

**Figure 3.**
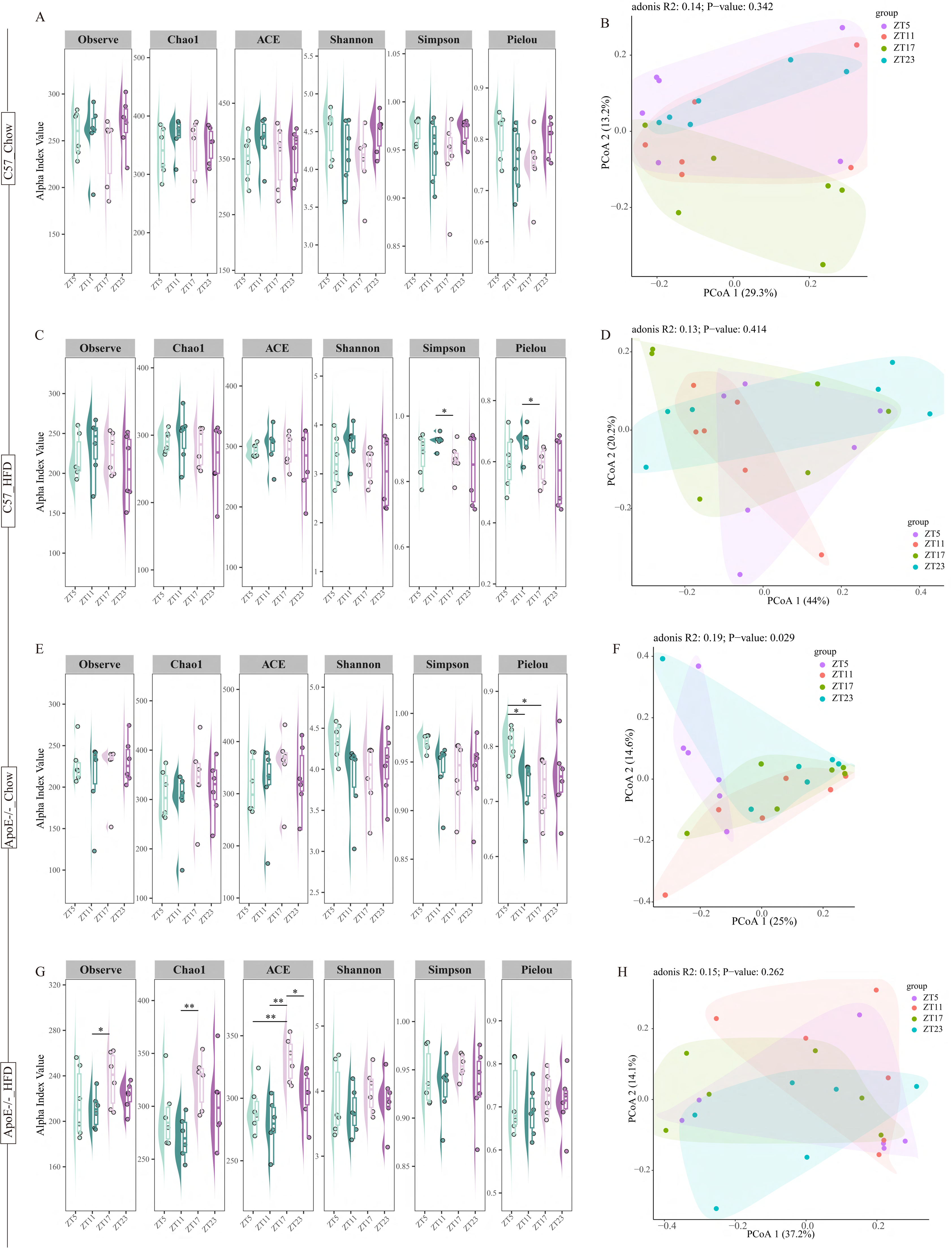
Dynamic change of α-diversity and β-diversity of fecal microbiota of C57_Chow, C57_HFD, ApoE-/-_Chow and ApoE-/-_HFD mice. (A, C, E, G) Dynamic change in α-diversity of fecal microbiota in C57_Chow, C57_HFD, ApoE-/-_Chow and ApoE-/-_HFD mice; (B, D, F, H) Dynamic change in β-diversity of fecal microbiota in C57_Chow, C57_HFD, ApoE-/-_Chow and ApoE-/-_HFD mice; Relative variable importance (R2) and significance (*P*) were calculated by PERMANOVA (Adonis) analysis. * indicates *P* < 0.05, ** indicates *P* < 0.01; n=6.

**Figure 4.**
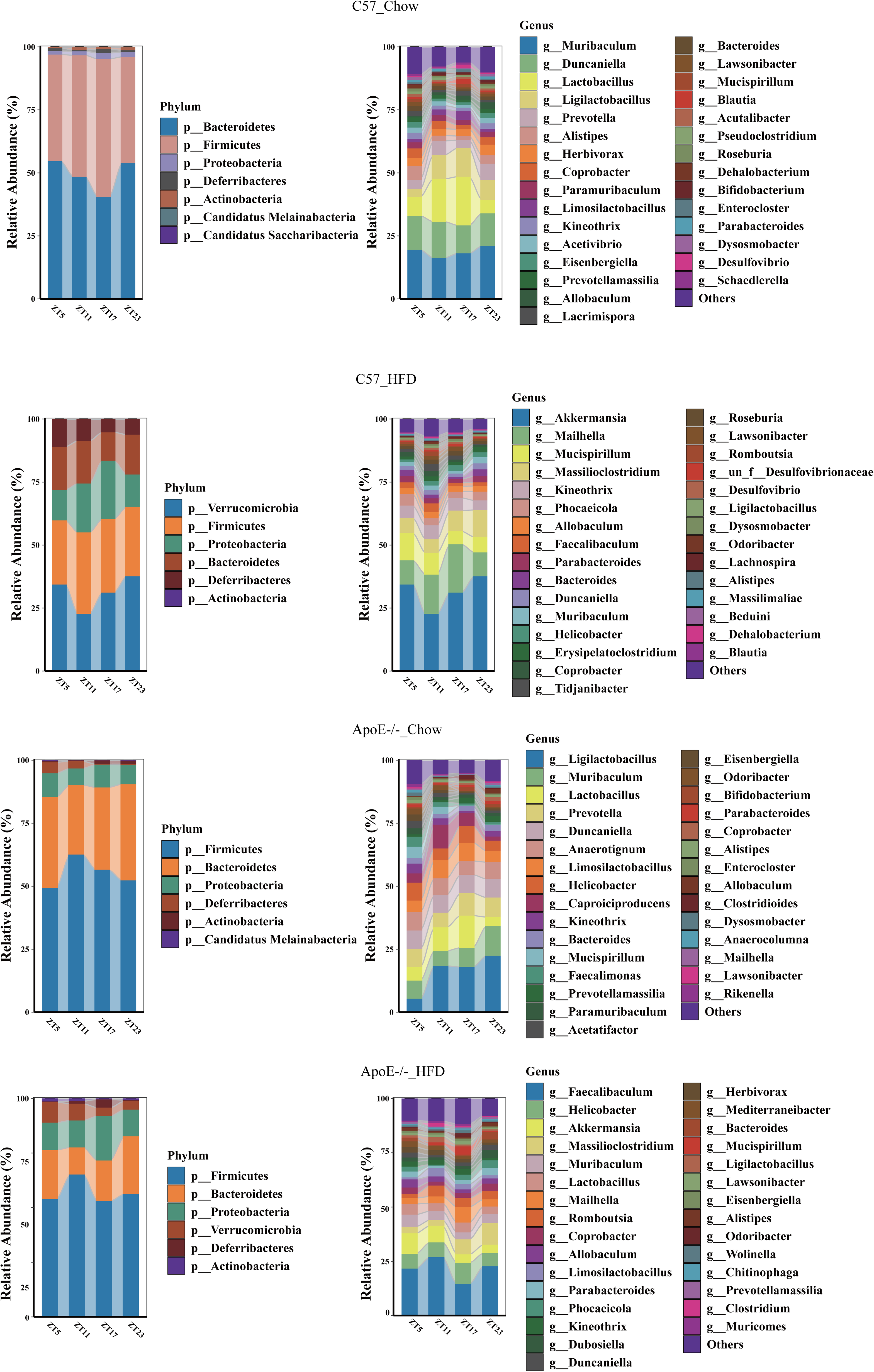
Dynamic change in fecal microbiota composition at the phylum and genus level of C57_Chow, C57_HFD, ApoE-/-_Chow, and ApoE-/-_HFD mice. (A, C, E, G) Dynamic change in fecal microbiota composition at the phylum in C57_Chow, C57_HFD, ApoE-/-_Chow and ApoE-/-_HFD mice; (B, D, F, H) Dynamic change in fecal microbiota composition at the genus in C57_Chow, C57_HFD, ApoE-/-_Chow and ApoE-/-_HFD mice. n=6.

**Figure 5.**
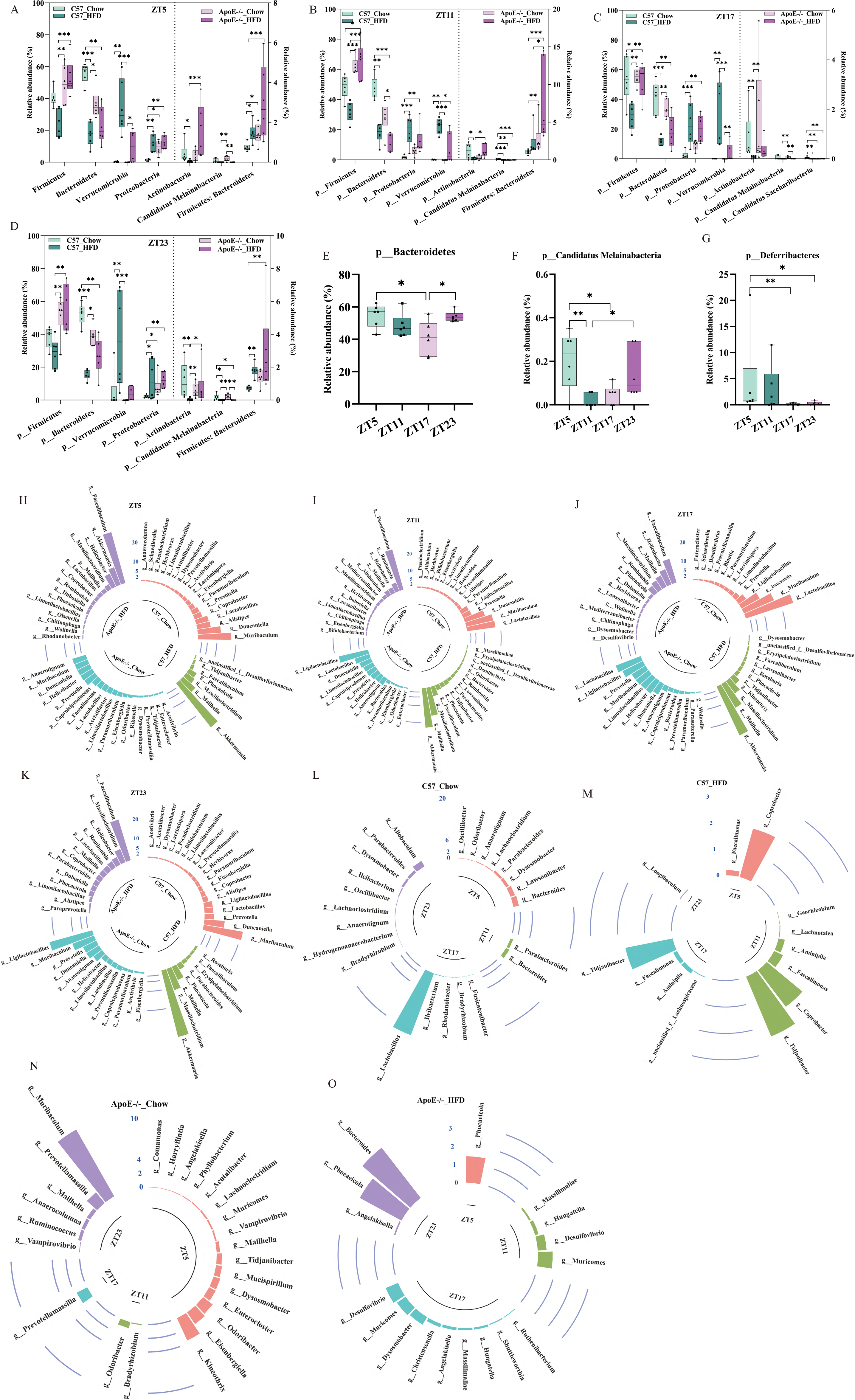
Differences in the relative abundance of fecal microbial at the phylum and genus level of C57_Chow, C57_HFD, ApoE-/-_Chow, and ApoE-/-_HFD mice. (A-D) Differences in the relative abundance of fecal microbiota at the phylum level among four groups of mice at ZT5, ZT11, ZT17, and ZT23, respectively. The relative abundance of each bacterial taxon is expressed as mean ± standard error. (E) Differences in relative abundance at the phylum level of fecal microbiota in C57_Chow mice at different time points. The relative abundance of each bacterial taxon is expressed as mean ± standard error. (F, G) Differences in relative abundance at the phylum level of fecal microbiota in ApoE-/-_Chow mice at different time points. The relative abundance of each bacterial taxon is expressed as mean ± standard error. (H-K) Differences in the relative abundance of fecal microbiota at the genus level among four groups of mice at ZT5, ZT11, ZT17, and ZT23, respectively (relative abundance>0.5%). Relative abundance levels are expressed as median; (L-O) Differences in the relative abundance of fecal microbiota at the genus level at different time points in C57_Chow, C57_HFD, ApoE-/-_Chow, and ApoE-/-_HFD, respectively. Relative abundance levels are expressed as median. * indicates *P* < 0.05, ** indicates *P* < 0.01, *** indicates *P* < 0.001; n=6.

At the phylum level, non-parameter JTK circle analysis showed significant oscillation in the phyla Firmicutes, Bacteroidetes, and F/B (Firmicutes: Bacteroidetes) of the fecal microbiota in C57_Chow group mice (Fig 6A-6C). Phyla Candidatus Melainabacteria and Deferribacteres of the fecal microbiota in ApoE^-/-^_Chow group mice underwent significant oscillation (Fig 6D, 6E). Further analysis revealed that there were significant differences in the genus and species that exhibited significant oscillation in the fecal microbiota of the four groups of mice (Fig 6F, 6G). In addition, non-parametric JTK analysis of microbiota function revealed significant differences in fecal microbial function among four groups of mice that exhibited significant oscillations (Fig S3). Compared with normal mice, the diurnal oscillatory rhythms of the composition and function of fecal microbiota in the mouse model of atherosclerosis changed significantly.

**Figure 6.**
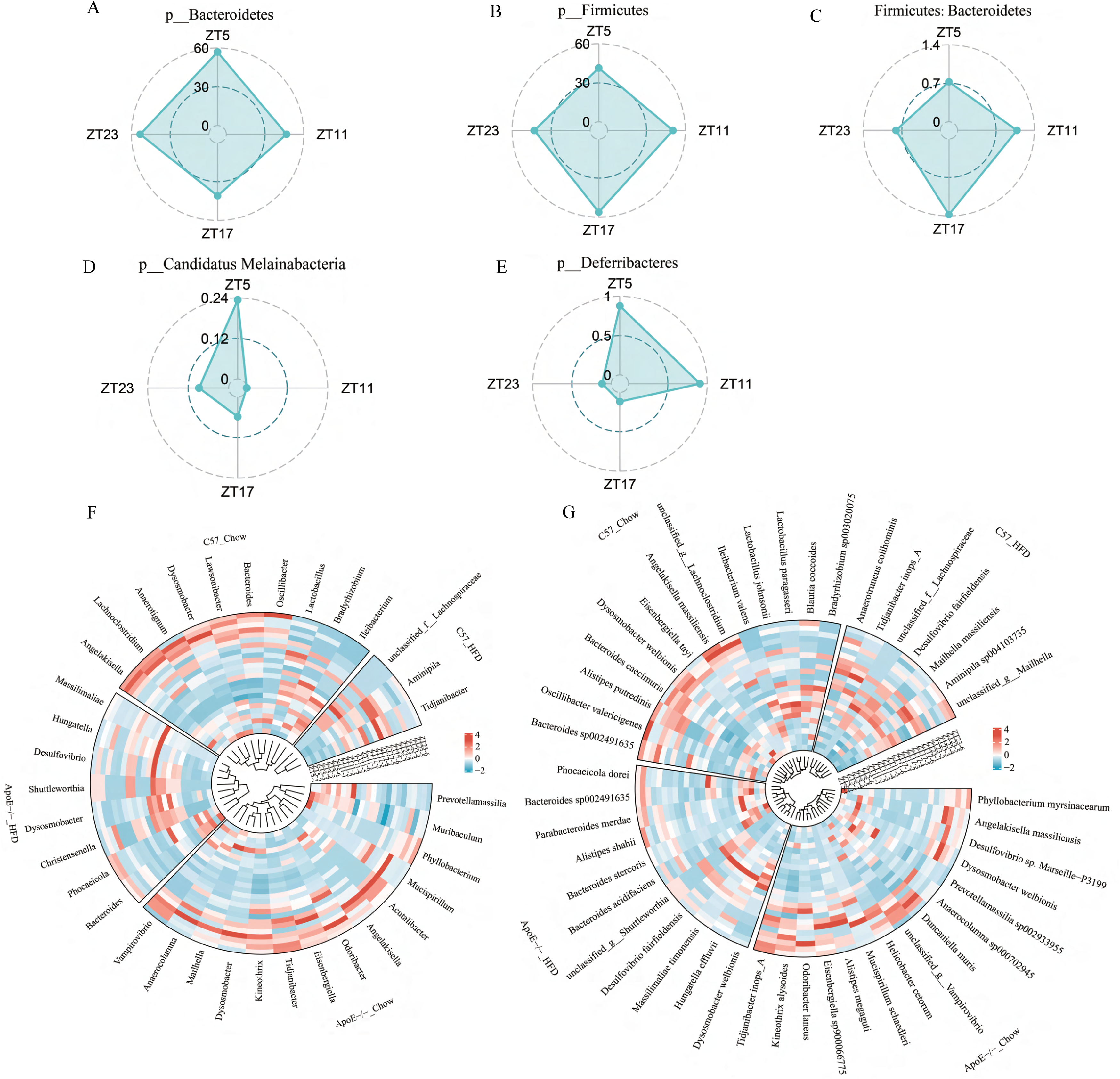
Daily fluctuation of the fecal microbiota at the phylum and genus level of C57_Chow, C57_HFD, ApoE-/-_Chow, and ApoE-/-_HFD mice. (A-C) Dynamic change of fecal microbiota at the phylum level of C57_Chow mice at each sampling time point within 24 h. Relative abundance levels are expressed as median. (D, E) Dynamic change of fecal microbiota at the phylum level of ApoE-/-_Chow mice at each sampling time point within 24 h. Relative abundance levels are expressed as median. (F, G) Dynamic change of fecal microbiota at the genus and species level of C57_Chow, C57_HFD, ApoE-/-_Chow, and ApoE-/-_HFD mice at each sampling time point within 24 h. The rhythmicity analysis was finished by non-parametric JTK analysis. n=6.

### Rhythm oscillation *Blautia coccoides* was closely related to the progression of ASCVD

The differential analysis of the relative abundance of microbial function (cardiovascular disease) in four groups of mice at different time points within a day found that the relative abundance of cardiovascular disease in ApoE^-/-^_HFD mice was significantly higher in C57BL/6 mice at ZT5, ZT17, and ZT23, except for ZT11 (Fig 7A). Moreover, the relative abundance of cardiovascular disease in ApoE^-/-^_Chow mice was significantly higher in C57_Chow mice at ZT17. To investigate the impact of rhythmic oscillatory strains on the progression of ASCVD, we employed LEfSe to analyze the fecal microbiota of the four groups of mice at ZT17 (Fig 7C). To further identify the key bacterial taxa associated with the progression of ASCVD, we regressed the abundance of LEfSe-distinguished bacterial taxa (Top 30) against the relative abundance of microbial function (cardiovascular disease) using the random forests machine learning algorithm (Fig 7E). We noticed that of the seven key bacterial taxa associated with the progression of ASCVD, only the relative abundance of *Blautia coccoides* was negatively correlated with the progression of AS (Fig 7E, S5). Interestingly, the relative abundance of *Blautia coccoides* in C57_Chow mice exhibited significant rhythmic fluctuations with a peak at ZT17 and a trough at ZT11 (Fig 7F). These results revealed that circadian rhythm dysfunction of gut microbiota is associated with the progression of ASCVD.

**Figure 7.**
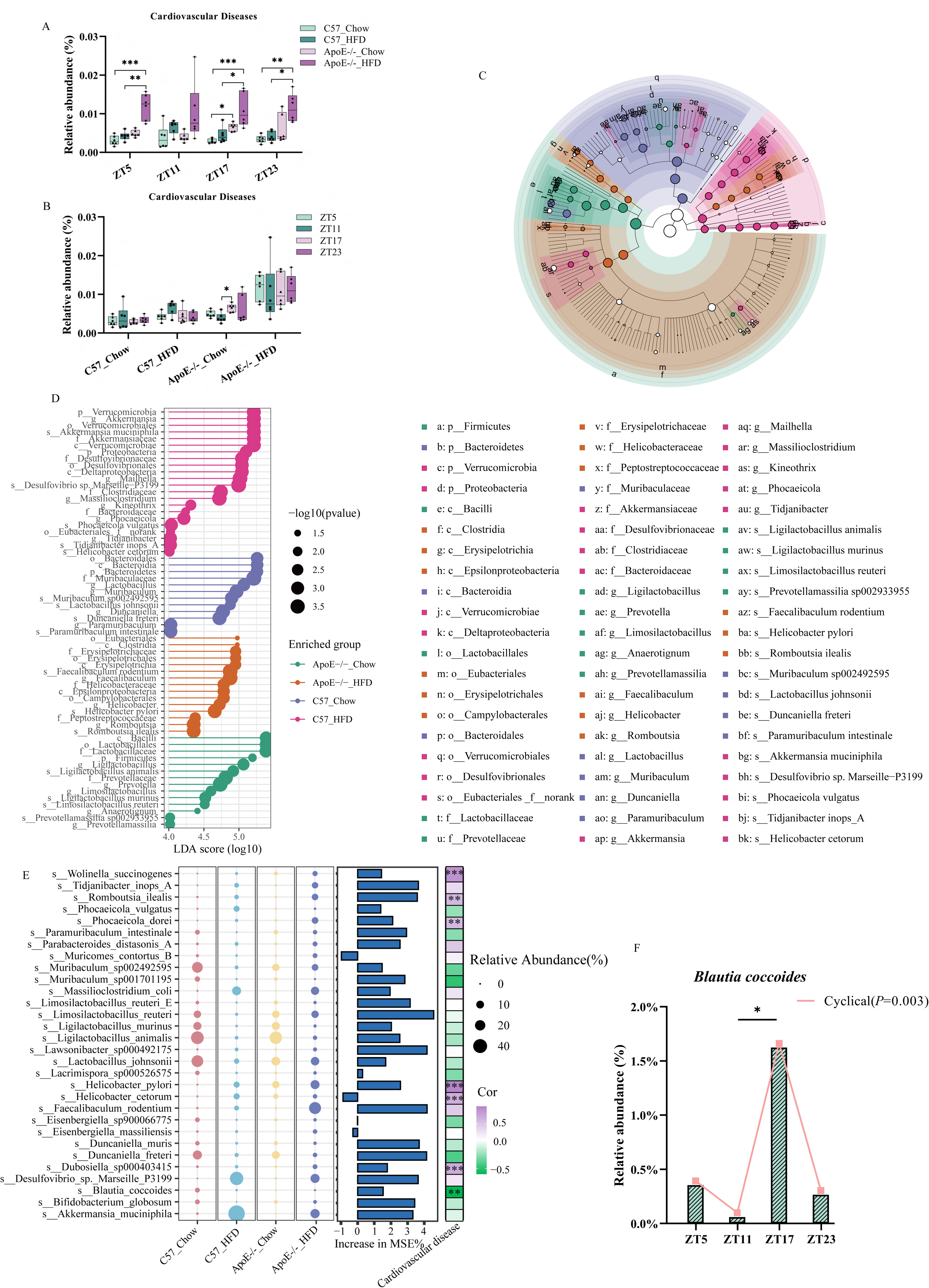
*Blautia coccoides* was closely related to cardiovascular disease. (A) Differences in the relative abundance of microbial function (Cardiovascular Diseases) among C57_Chow, C57_HFD, ApoE-/-_Chow, and ApoE-/-_HFD mice at ZT5, ZT11, ZT17, and ZT23, respectively. Relative abundance was expressed as mean ± standard error. (B) Differences in the relative abundance of microbial function (Cardiovascular Diseases) among ZT5, ZT11, ZT17, and ZT23 of C57_Chow, C57_HFD, ApoE-/-_Chow and ApoE-/-_HFD mice, respectively. Relative abundance was expressed as mean ± standard error. (C, D) Linear discriminant analysis Effect Size (LEfSe) analysis of the microbiota among C57_Chow, C57_HFD, ApoE-/-_Chow, and ApoE-/-_HFD mice (LDA > 4, *P* < 0.01). (E) The top 7 bacterial biomarkers were identified by random forests regression of relative abundances of LEfSe-determined bacterial taxon (Top 30 species) against microbial function (Cardiovascular Diseases). The statistical significance of selected bacterial biomarkers was assessed by permutation test (999 times). (F) Diurnal oscillation of relative abundance of *Blautia coccoides* in fecal microbiota of C57_Chow group mice. * indicates *P* < 0.05, ** indicates *P* < 0.01, *** indicates *P* < 0.001; n=6.

### Oral gavage with *Blautia coccoides* protected against HFHCD diet-induced atherosclerotic lesion formation in ApoE-/- mice

We next investigated whether *Blautia coccoides* could alleviate the progression of AS. 5-week-old male ApoE^-/-^ mice were administrated with either PBS, heat-killed *Blautia coccoides* (Ina-B.Coccides), or live *Blautia coccoides* (B. Coccides) by oral gavage for 11 weeks after 2-week of interference with broad-spectrum antibiotics. As expected, oral administration of live *Blautia coccoides* to HFHCD mice substantially enhanced the abundance of *Blautia coccoides* in the murine feces, compared to that in PBS or Ina-B.Coccides treated HFHCD mice (Fig. 8A). We further observed that *Blautia coccoides* intervention significantly improved growth performance, as shown by reduced body weight and the relative weight of epididymal adipose tissue (EAT), compared to that in PBS or Ina-B.Coccides treated HFHCD mice (Fig. 8B-8D). In addition, *Blautia coccoides* intervention did not exhibit a significant difference in plasma CHOL, TG, and HDL-C concentration, whereas the plasma LDL-C concentration was decreased (Fig. 8E-8H).

**Figure 8.**
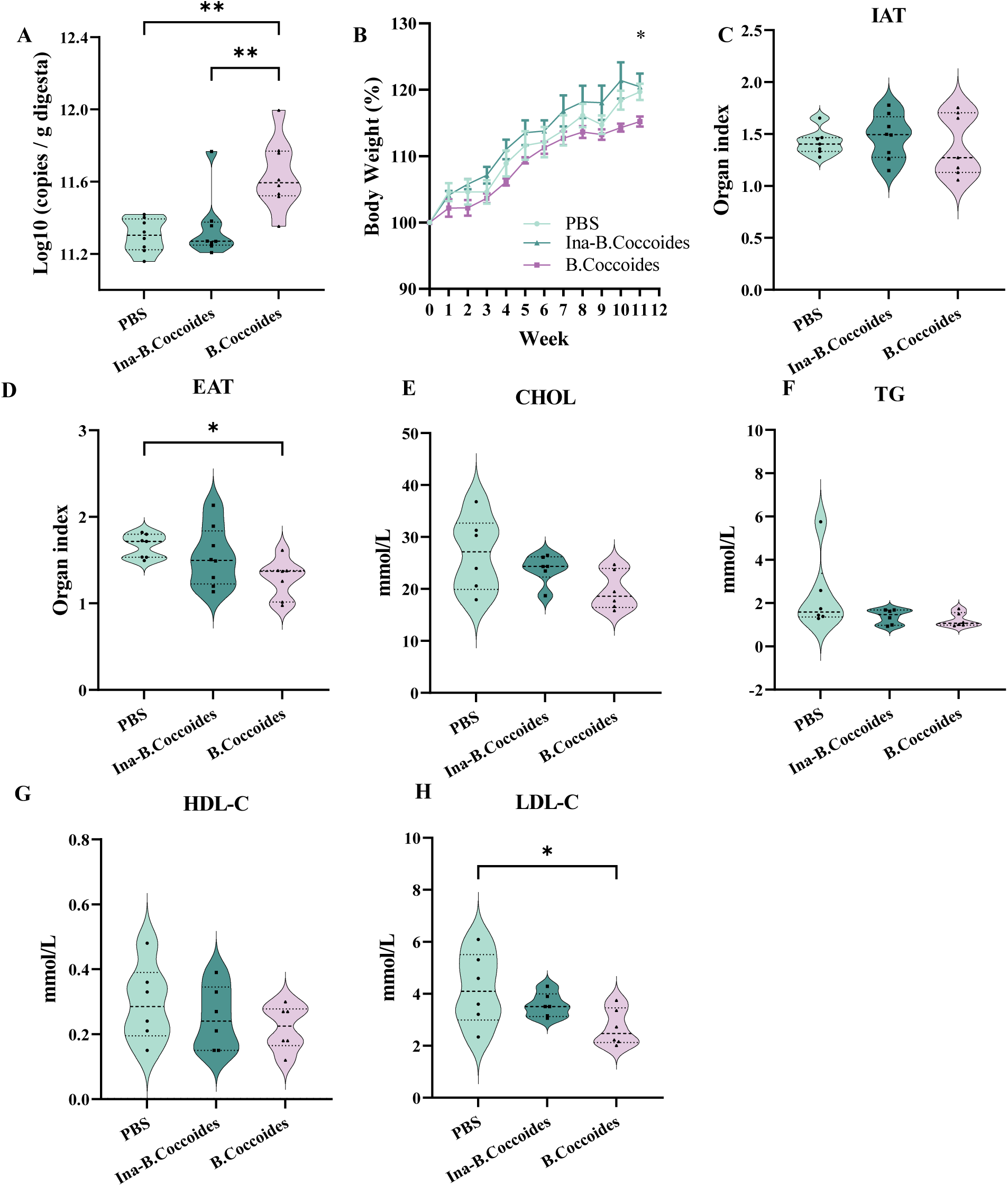
*Blautia coccoides* intervention influences body weight gain and blood lipid level in ApoE mice. (A) Absolute abundance of *Blautia coccoides* in colonic contents after 11 weeks of intervention. (B) Mice were maintained on HFD with oral administration of PBS, heat-killed *Blautia coccoides* (Ina-B. Coccides), or live *Blautia coccoides* (B. Coccides) three times each week and were weighed weekly. (C, D) Effect of *Blautia coccoides* intervention on the relative weight of subcutaneous inguinal adipose tissue (IAT) and epididymal adipose tissue (EAT). (E-H) Effect of *Blautia coccoides* intervention on blood lipid level. Data was expressed as mean ± standard error. *P* < 0.05, ** indicates *P* < 0.01; n=6.

The *Blautia coccoides* administrated protected against HFHCD diet-induced atherosclerotic lesion formation, as demonstrated by Oil Red O staining of entire aorta and aortic root regions, and hematoxylin and eosin staining of aortic root regions. Quantification of atherosclerotic plaque as measured in the entire aorta stained using Oil Red O, showed that the *Blautia coccoides* administrated significantly decreased the lesion area in ApoE^-/-^mice compared with PBS or Ina-B.Coccides group mice (Fig 9A). In addition, the results of plaque area as measured in aortic roots (cross-section method) also showed that *Blautia coccoides* administrated significantly reduced plaque accumulation compared with PBS or Ina-B.Coccides group mice (Fig 9C). Administration of the same dose of heat-killed *Blautia coccoides* did not show any improvement in lesion area or size in ApoE^-/-^mice, indicating that the protective effect of these bacteria was dependent on their viability.

**Figure 9.**
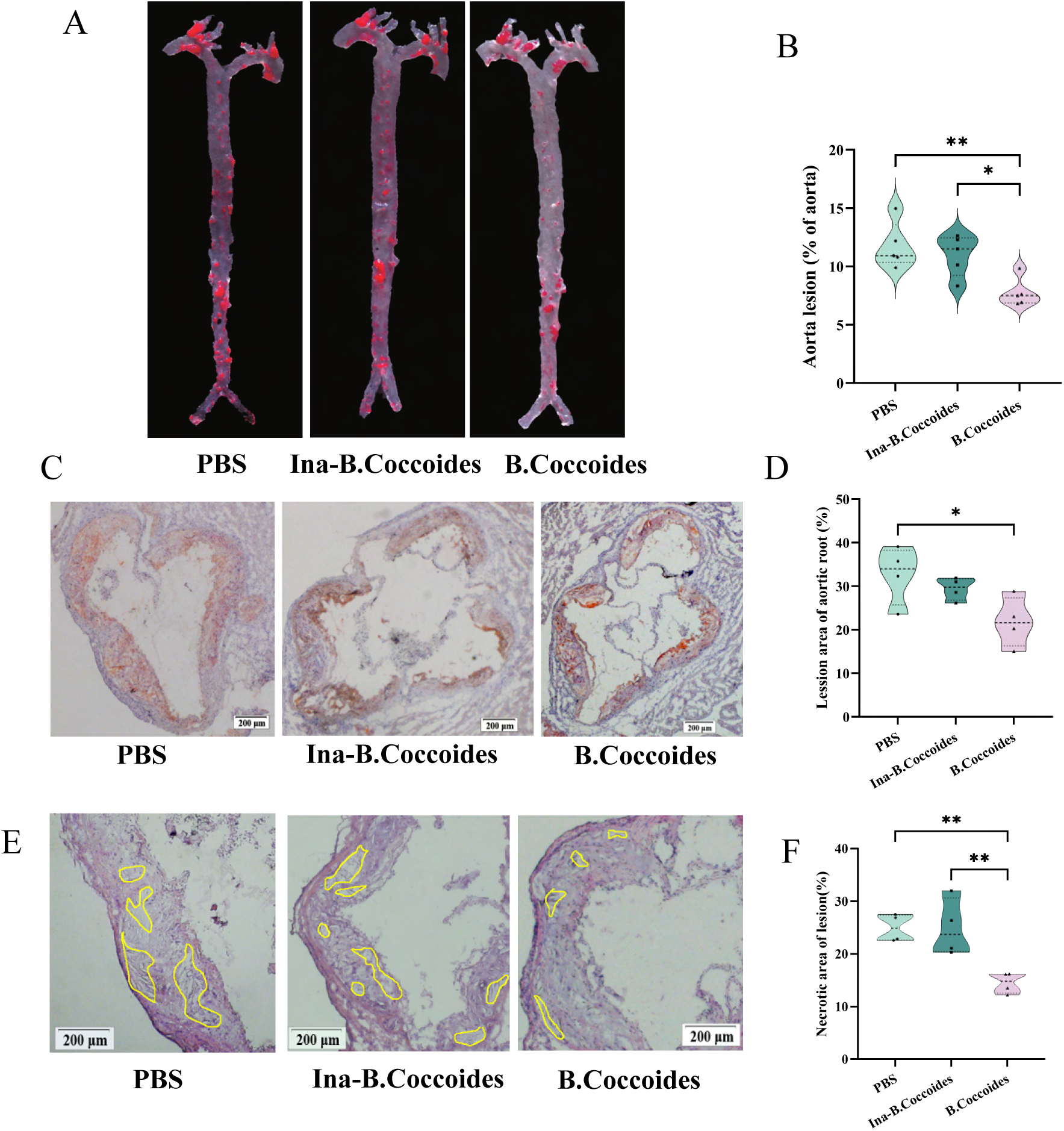
*Blautia coccoides* intervention inhibits the progression of atherosclerosis. (A, B) Representative photomicrographs of oil red O staining and quantitative analysis of atherosclerotic plaque area in the whole aortas. (C, D) Representative oil red O and quantification of the aortic sinus lesion area. (E, F) Representative hematoxylin and eosin staining and quantification of the aortic sinus lesion area. Data was expressed as mean ± standard error. *P* < 0.05, ** indicates *P* < 0.01; n=4.

### Metabolites from *Blautia coccoides* protected against HFHCD diet-induced atherosclerotic lesion formation by improved endothelial barrier function

To identify potent *Blautia coccoides*-derived metabolites that exhibit beneficial effects in protecting against atherosclerotic lesion formation induced by HFHCD, we performed non-targeted metabolome sequencing of the supernatant from *Blautia coccoides* by using LC-MS/MS. Data analysis of the resulting volcano plot revealed the gallic acid (GA) was increased more than 52-fold in the supernatant of *Blautia coccoides* cultures compared with uncultured, fresh medium (Fig. 10A). We treated MAEC cells with different concentrations of GA to explore the potential mechanism of GA on the progression of ASCVD in vitro. The qPCR results showed that different concentrations of GA had no significant effect on the expression of genes (e.g. *ABCA1*, *SR-B1*, and *APOA1*) mediating reverse cholesterol transport (Fig. 10B-10D). The relative expression level of TNFα was significantly decreased by GA (2 μg/mL). However, GA did not significantly affect the relative expression levels of genes (e.g. *NLRP3*, *MCP1*, and *IL6*) involved in inflammation response (Fig. 10E-10H). In addition, GA treatment significantly increased the relative expression of VE-cadherin in MAEC (Fig. 10I). Together, GA can inhibit the inflammatory response of MAEC and improve endothelial barrier function.

**Figure 10.**
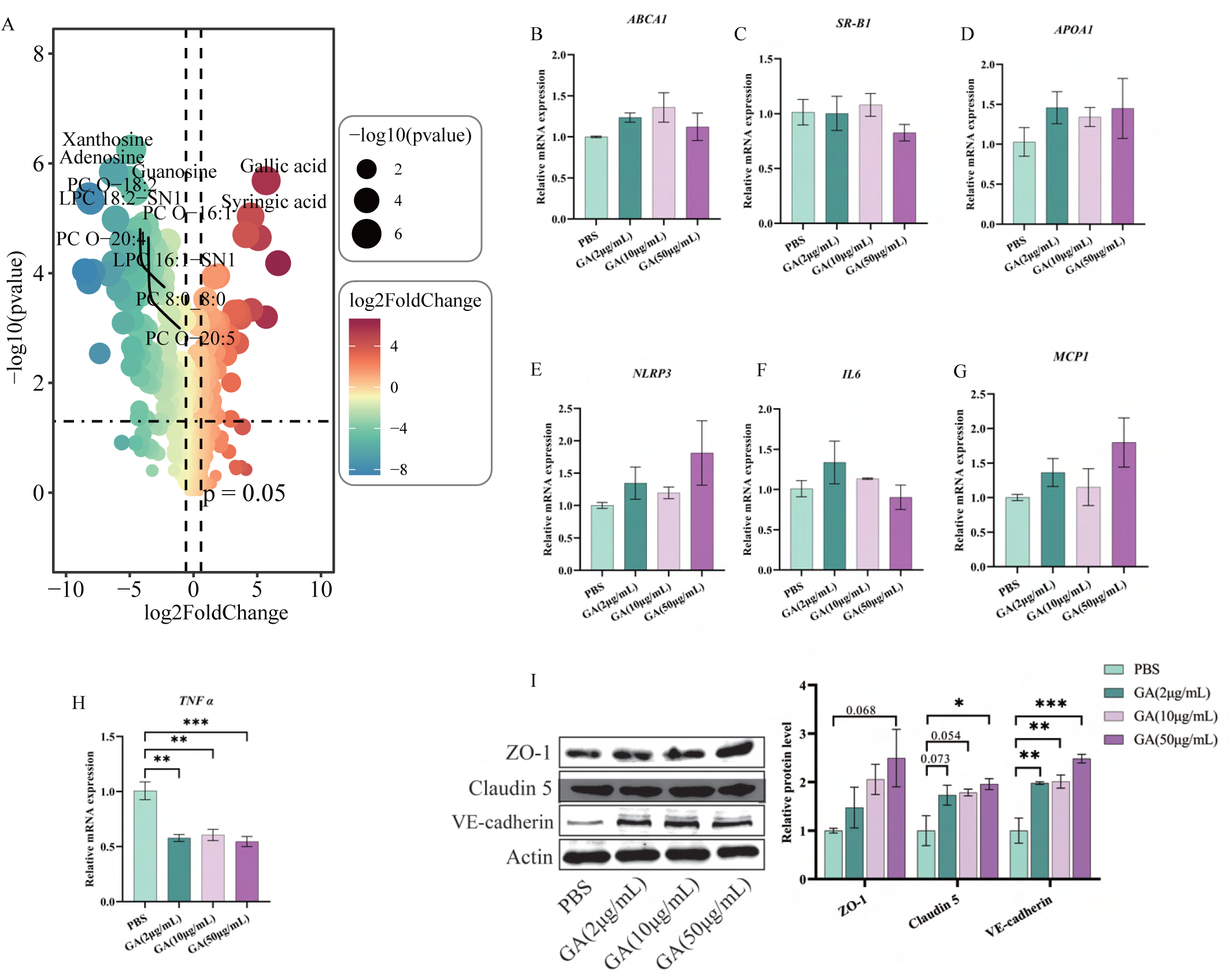
Metabolites from *Blautia coccoides* protected against HFHCD diet-induced atherosclerotic lesion formation. (A) Volcano plot showing LC–MS/MS analysis of *Blautia coccoides* culture supernatant. Red and blue dots indicate metabolites increased and decreased, respectively (n = 3). (B-D) The effects of GA on cholesterol transport in mouse aortic endothelial cells (MAEC). (n=3). (E-H) The effects of GA on the inflammatory response of MAEC. (n=3). (I) The effects of GA on the endothelial barrier function of MAEC. (n=3). Data was expressed as mean ± standard error. * indicates *P* < 0.05, ** indicates *P* < 0.01, *** indicates *P* < 0.001, numbers indicates *P*-values. GA, gallic acid.

## Discussion

Recent advancements in microbiome research have highlighted a robust association between gut dysbiosis and the progression of ASCVD^[8, 9, 21–23]^, supporting that metabolite-derived gut microbiota can function alone through pathways involved in host metabolism, contributing to health and disease processes. However, the intricate interplay between diet, lifestyle, gut microbiota, and the host significantly impacts the progression of ASCVD, leading to the precise mechanisms of which and how specific gut microbial metabolites contribute to the progression of atherosclerosis, and the clinical relevance of their alterations remains unclear. Here, our data suggested that the diurnal oscillatory rhythms of fecal microbiota composition and function in ApoE^-/-^ mice are significantly altered. Furthermore, we demonstrated that the relative abundance of the rhythmic strain (*Blautia coccoide*s) was significantly negatively correlated with the progression of atherosclerosis, and its metabolites (GA) could protect against atherosclerotic lesion formation by improving the function of the endothelial barrier. Our study supports that circadian rhythm plays an important role in the progression of ASCVD^[24–27]^, and provides new insights into the mechanisms by which dietary behavior regulates the progression of ASCVD.

Lifestyle and nutritional habits serve as pivotal factors in triggering immune-metabolic disorders, including ASCVD ^[28, 29]^. Following healthy eating habits, such as a deliberate daily pause in food consumption, maybe a potentially cost-effective strategy for reducing the risk of chronic diseases ^[30]^. However, due to the diversity of dietary patterns and the limitations of research, the potential regulatory mechanisms by which dietary habits affect the development of ASCVD remain unclear. Changes in eating habits have been shown to improve immune-metabolic functions by sharpening their circadian rhythm. Short-term circadian misalignment can increase 24-h blood pressure and inflammatory markers in healthy adults, which may help explain why circadian misalignment is a risk factor for cardiovascular disease ^[25]^. The gut microbiota is closely associated with ASCVD and exhibits circadian oscillations in composition and function ^[19]^. However, whether diurnal oscillatory rhythms of the gut microbiota are involved in regulating the progression of atherosclerosis remains unclear.

In line with the previous study, the total numbers of cycling 16S rRNA gene amplicon sequence variants in C57BL/6 mice with ad libitum HCHFD feeding was more severely reduced than that in C57BL/6 mice with ad libitum chow-fed groups ^[31]^. In addition, our findings revealed that the HCHFD also reduced the number of operational taxonomic units (OTUs) in ApoE-/- mice (Fig. S2). Compared with C57_Chow mice, HCHFD reduced the relative abundance of Firmicutes in feces. However, the relative abundance of Firmicutes was the highest in ApoE^-/-^ mice, regardless of standard diet or HFHCD. To characterize the cyclical oscillations of the fecal microbiome and determine their importance in host metabolism, we investigated the relative abundances of taxa under different diets over time. Through the analysis of diurnal oscillations in microbiota composition using the nonparametric algorithm JTK_cycle, we found that the C57BL/6 and ApoE^-/-^ mice in the ad libitum HCHFD feeding group exhibited fewer diurnal oscillatory rhythms at the phylum, genus, and species level than mice in the ad libitum chow-fed groups.

Changes in the diurnal oscillatory pattern of microbial composition significantly affected the diurnal oscillatory of fecal microbiota functions. There were significant differences in the diurnal cycle of the relative abundance of carbohydrate metabolism (e.g. Amino sugar and nucleotide sugar metabolism, Fructose and mannose metabolism, and Inositol phosphate metabolism) and lipid metabolism (e.g. Secondary bile acid biosynthesis, Primary bile acid biosynthesis) in C57BL/6 mice, regardless standard diet or HFHCD (Fig S4). However, no differences were found in the diurnal cycle of carbohydrate metabolism and lipid metabolism in ApoE^-/-^ mice. The ability of gut microbiota to reduce cholesterol by modifying intestinal bile acids is considered an important dietary intervention for reducing disease risk and improving population health ^[32, 33]^. Meanwhile, bile acids have also provided a link between gut microbiota and cardiovascular health. Changes in lipid metabolism of microbiota function in ApoE^-/-^ mice, specifically on bile acid biosynthesis, may be one of the reasons why diurnal oscillatory microbiota impacts the progression of ASCVD. Our findings suggested significant changes in the diurnal dynamics of the composition and function of ApoE^-/-^ mice fecal microbiome.

The genus *Blautia* in the gut microbiota of Japanese people is more abundant than that of people from other countries, and studies involving Japanese cohorts consistently indicate the beneficial effects of *Blautia* ^[34]^. Numerous studies have highlighted that *Blautia* is closely related to metabolism diseases ^[35]^. *Blautia coccoides* belongs to the *Clostridium* class, which is one of the major gut microbiotas in animals and humans, and has a significant impact on intestinal health ^[36, 37]^. Our research indicated that the relative abundance of *Blautia coccoides* in C57_Chow mice exhibited significant diurnal oscillatory and was negatively correlated with the progression of ASCVD. Circadian rhythm disturbances contribute to disruption of the gut microbiome, which may be by reducing gut barrier integrity and immune responses ^[38]^. *Blautia* has been identified as a mucus-associated bacterium in the colon of humans and mice ^[39, 40]^, and recent findings^[37]^ indicated that *Blautia coccoides* could serve as a new target to restore mucus growth during mucus-associated lifestyle diseases, a process that stimulated mucus growth through the production of the short-chain fatty acids (eg. propionate and acetate) via activation of the short-chain fatty acid receptor. Compared with the PBS or Ina-B.Coccides treated HFHCD mice, oral gavage with *Blautia coccoides* significantly decreased the relative expression level of NF-κB in the colon of ApoE^-/-^ mice (Fig. S6). In addition, compared with the PBS or Ina-B.Coccides treated HFHCD mice, oral gavage with *Blautia coccoides* significantly increased the relative expression level of Claudin-1 in the colon of ApoE^-/-^ mice. There was a notable tendency towards increased expression levels of ZO-1 protein (*P* = 0.07) in B.Coccides group. In line with previous studies, our findings indicated that *Blautia coccoides* can effectively improve the colonic barrier function in mice.

Metabolomics data showed that GA concentration in the supernatant of *Blautia coccoides* culture medium increased more than 52-fold than that in fresh medium. GA is a phenolic compound, often used as a dietary supplement or additive, and has been widely used against multiple conditions due to its antioxidant and anti-inflammatory properties ^[41–43]^. Our results indicate that GA could significantly reduce the levels of TNFα in MAEC, which was consistent with previous findings that GA could reduce the levels of TNFα in acute inflammation rats model ^[44]^. Endothelial dysfunction caused by the formation and release of inflammatory cytokines is a prominent feature of ASCVD vascular injury ^[45]^. TNF-α significantly reduced the expression of VE-cadherin, a vascular endothelial marker, which plays a central role in controlling endothelial barrier function ^[46]^. In endothelial cells, the vascular barrier composed of tight junctions and adhesive junctions plays a crucial role in maintaining endothelial integrity, effectively protecting the vessels from penetration and inflammation^[47, 48]^. Our results indicated that GA at a concentration of 2 μg/mL could significantly inhibit the expression of TNF and increase the expression of VE-cadherin, indicating that GA could improve endothelial barrier function.

## Conclusion

This study investigated a 24-h diurnal fluctuation of the composition and function in the fecal microbiota of C57BL/6 and ApoE^-/-^ mice with a standard chow diet or a high-fat, high-cholesterol diet under ad libitum conditions. We observed significant differences in the composition and function of the fecal microbiota in C57BL/6 mice and ApoE^-/-^ mice, as well as in their dynamic changes, regardless of whether they were fed a standard diet or a HFHCD. Random forest analysis found that rhythmic strain (*Blautia coccoides*) was significantly negatively associated with ASCVD progression. In addition, our results showed that *Blautia coccoides* inhibited ASCVD progression by improving the colon barrier and aortic endothelial cell barrier function. Together, our findings can provide a theoretical basis for understanding the role of diurnal rhythm of gut microbiota in regulating the progression of ASCVD, and simultaneously provide new ideas for the development of microbial-targeted therapeutic strategies for ASCVD.

## Ethical Approval

Animal Care and Use Committee of Xuzhou Medical University, Jiangsu, China (Code of Ethics: 202405T019) approved and supervised the experimental protocol for the care and treatment of mice in the study, and the animal experiments were conducted at the Experimental Animal Center of Xuzhou Medical University.

## Data sharing

The high-throughput datasets are available at NCBI project PRJNA1141708.

## Author contributions

He Zhang and Erteng Jia conceived and designed the study; He Zhang, Xiaohan Zhang, Zihan Yun, Yating Shao, Suhua Cang conducted the research; He Zhang, Erteng Jia, and Yang Chen analyzed and interpreted the data; He Zhang wrote the manuscript; Renjin Chen revised the manuscript. All authors read and approved the final version of the manuscript. The authors declare that all data were generated in-house and that no paper mill was used.

## Consent to Participate

Not applicable.

## Consent for Publication

All authors give their consent for publication of this manuscript.

## Acknowledgments

Not applicable.

---

**Figure S1.**
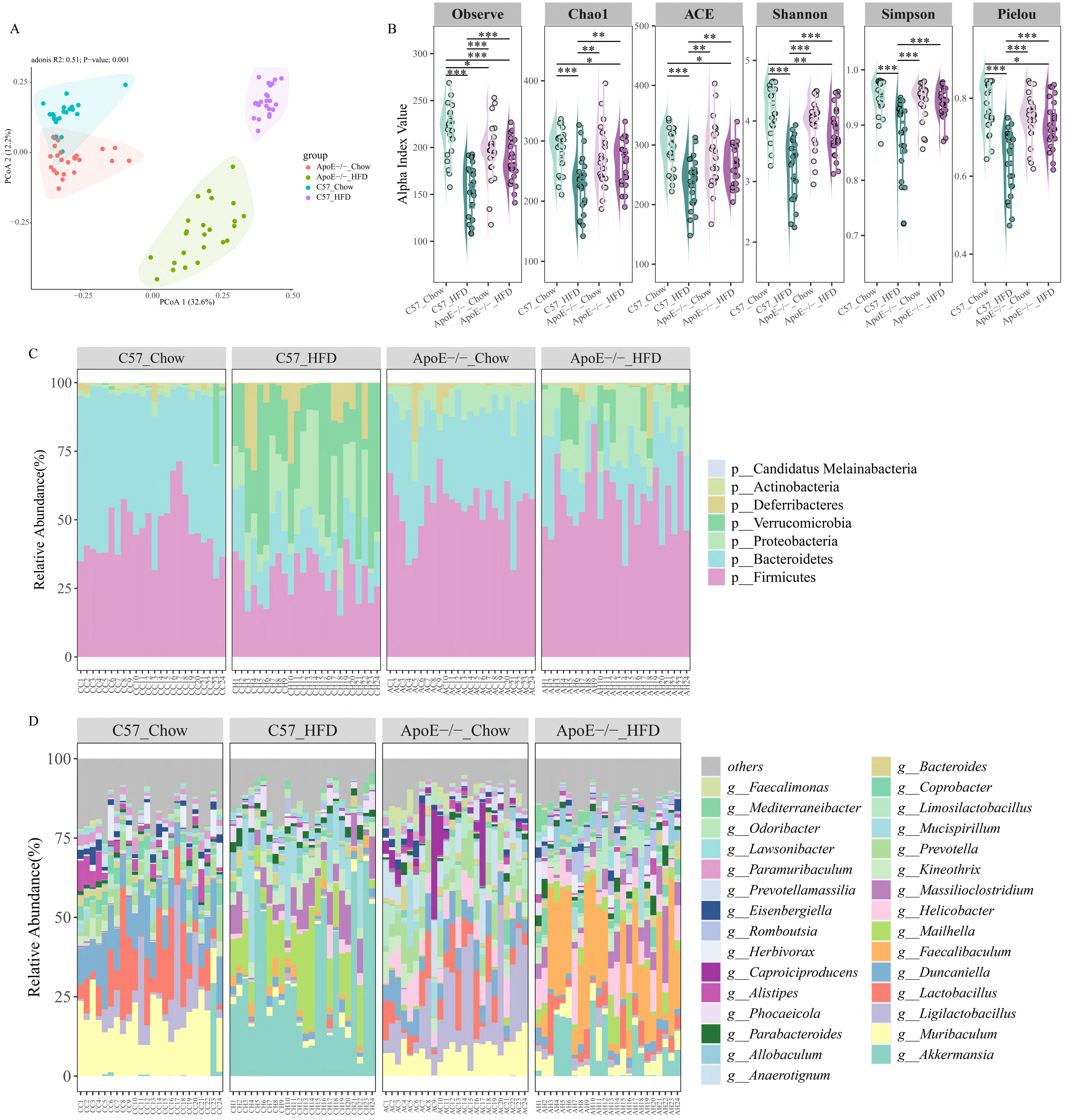
Fecal bacterial composition of C57_Chow, C57_HFD, ApoE-/-_Chow, and ApoE-/-_HFD mice under *ad libitum* conditions. (A) Principal-coordinates analysis (PCoA) of fecal microbiota among C57_Chow, C57_HFD, ApoE-/-_Chow, and ApoE-/-_HFD mice; PCoA was performed based on the Bray–Curtis metric. Relative variable importance (R2) and significance (*P*) were calculated by PERMANOVA (Adonis) analysis; (B) α-diversity of the fecal microbiota among C57_Chow, C57_HFD, ApoE-/-_Chow and ApoE-/-_HFD mice; (C) Barplot of fecal microbiota at the phylum level. Each bar represents an individual mouse and each color an individual phylum; (D) Barplot of fecal microbiota at the genus level. Each bar represents an individual mouse and each color is an individual genus. C57_Chow, C57BL/6 mice with standard diet; C57_HFD, C57BL/6 mice with high-fat, high-cholesterol diet; ApoE-/-_Chow, ApoE-/- mice with standard diet; ApoE-/-_HFD, ApoE-/- mice with high-fat, high-cholesterol diet. * indicates *P* < 0.05, ** indicates *P* < 0.01, *** indicates *P* < 0.001; n=24.

**Figure S2.**
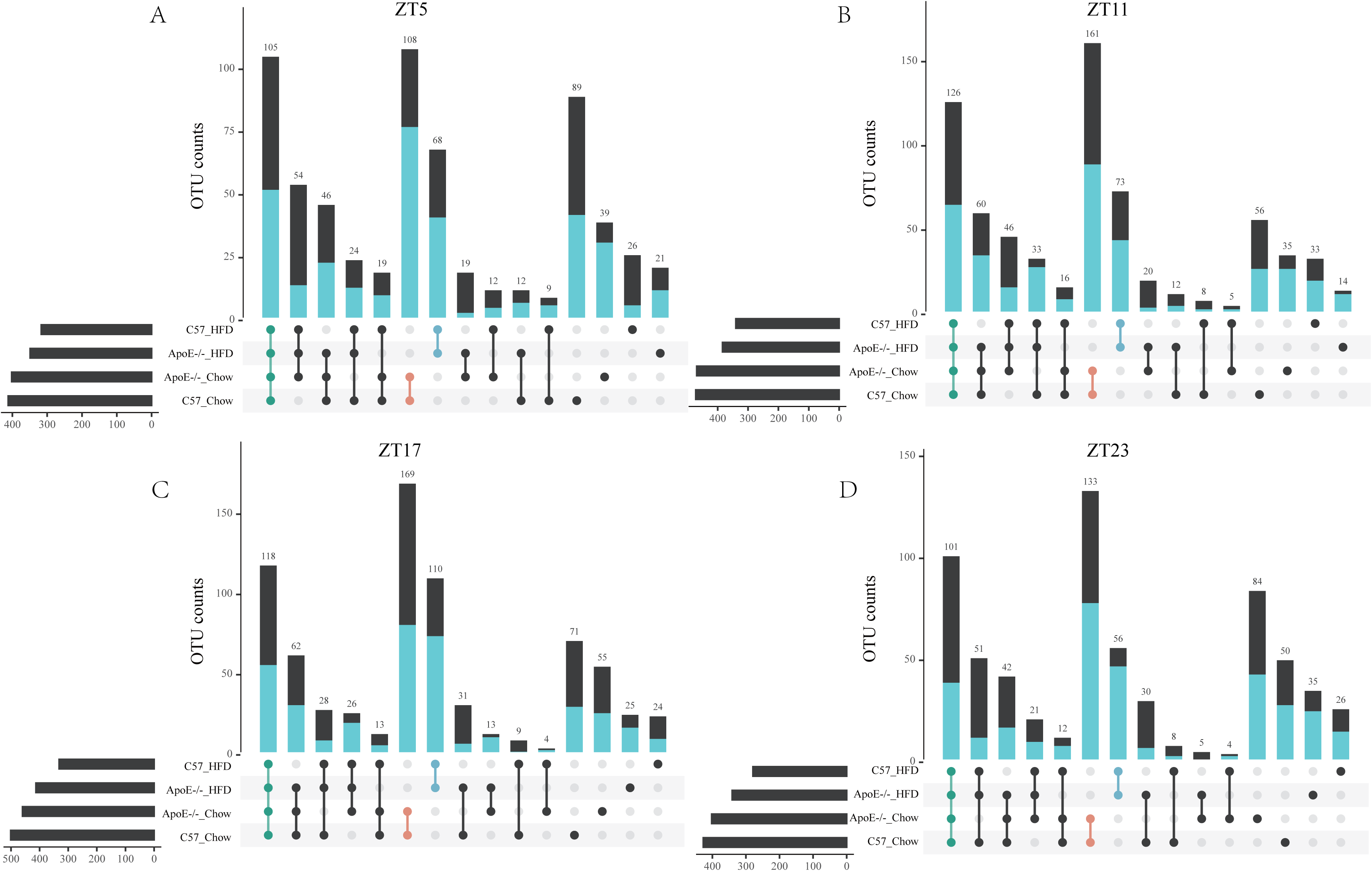
OTU counts of fecal microbiota in C57_Chow, C57_HFD, ApoE-/-_Chow, and ApoE-/-_HFD mice at different time points within a day. (A) ZT5; (B) ZT11; (C) ZT17; (D) ZT23. Filter out OTUs within the group that have an OTU count of 0 for more than half of their occurrences within the same group. The cyan columns represent OTU belonging to Firmicutes.

**Figure S3.**
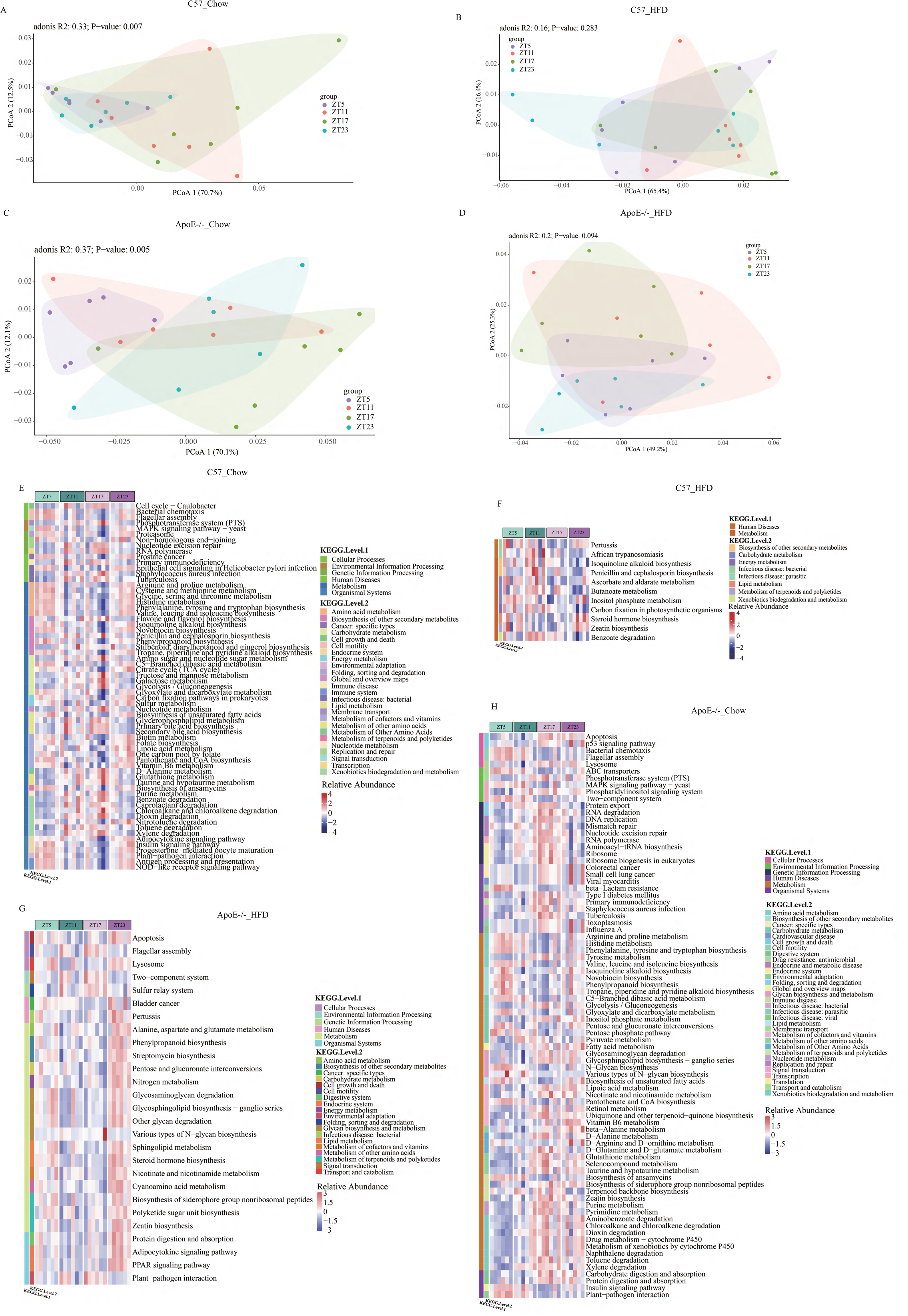
Daily fluctuation of microbial functions of C57_Chow, C57_HFD, ApoE-/-_Chow, and ApoE-/-_HFD mice. (A-D) PCoA of fecal microbial function at ZT5, ZT11, ZT17, and ZT23. Relative variable importance (R2) and significance (P) were calculated by PERMANOVA (Adonis) analysis. (E-H) Heatmap depicting the relative abundance of rhythmic KEGG level3 of fecal microbial function C57_Chow, C57_HFD, ApoE-/-_Chow, and ApoE-/-_HFD, respectively. PICRUSt2 was used to predict changes in microbial functions that might be associated with changes in OTU abundance detected through 16S sequencing. The rhythmicity analysis was finished by non-parametric JTK analysis. n=6.

**Figure S4.**
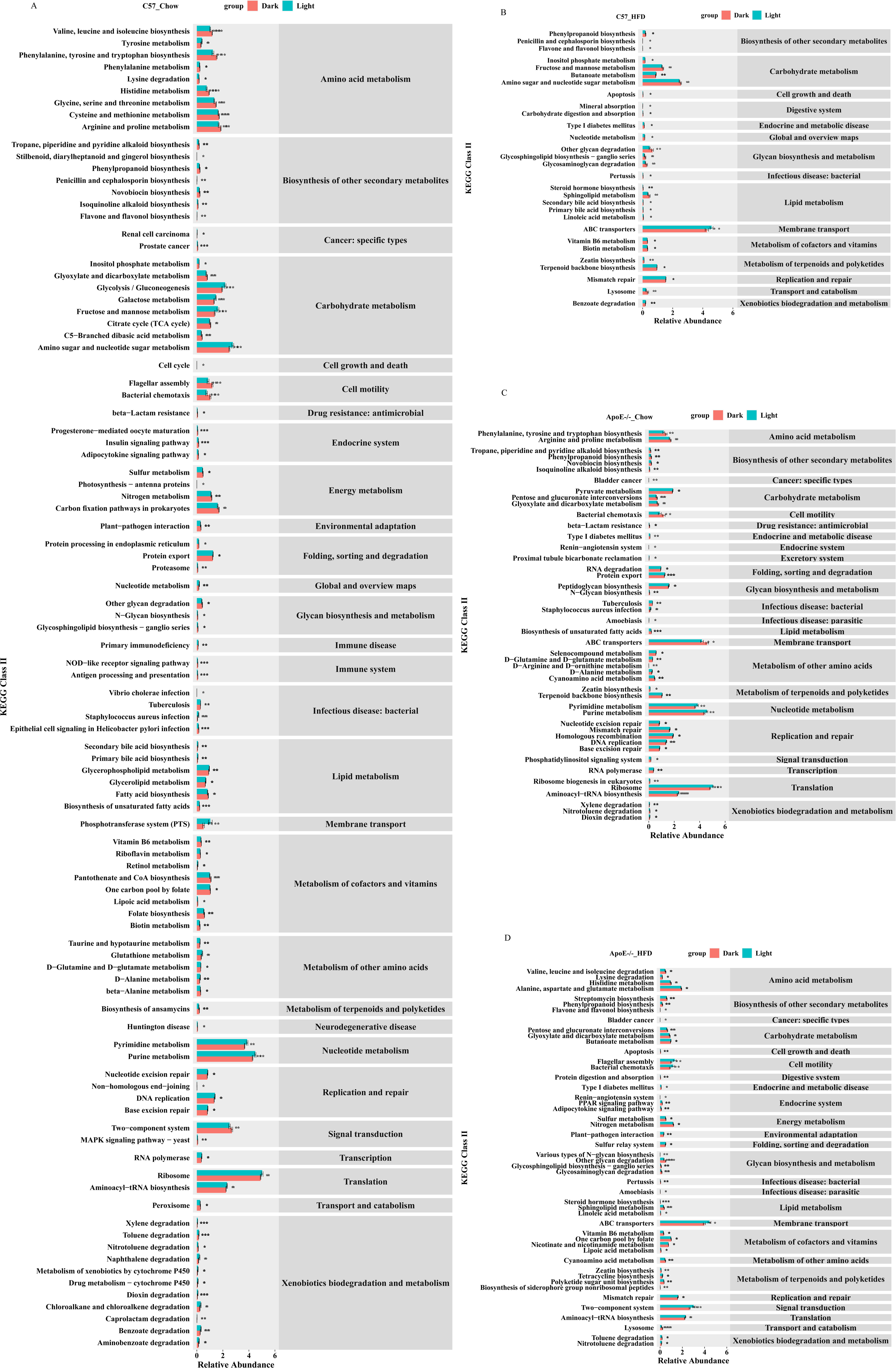
Daily differences in microbial functions of C57_Chow, C57_HFD, ApoE-/-_Chow, and ApoE-/-_HFD mice. (A) C57_Chow mice; (B) C57_HFD mice; (C) ApoE^-/-^_Chow mice; (D) ApoE^-/-^ HFD mice. Relative abundance was expressed as mean ± standard error. * indicates *P* < 0.05, ** indicates *P* < 0.01, *** indicates *P* < 0.001; n=12.

**Figure S5.**
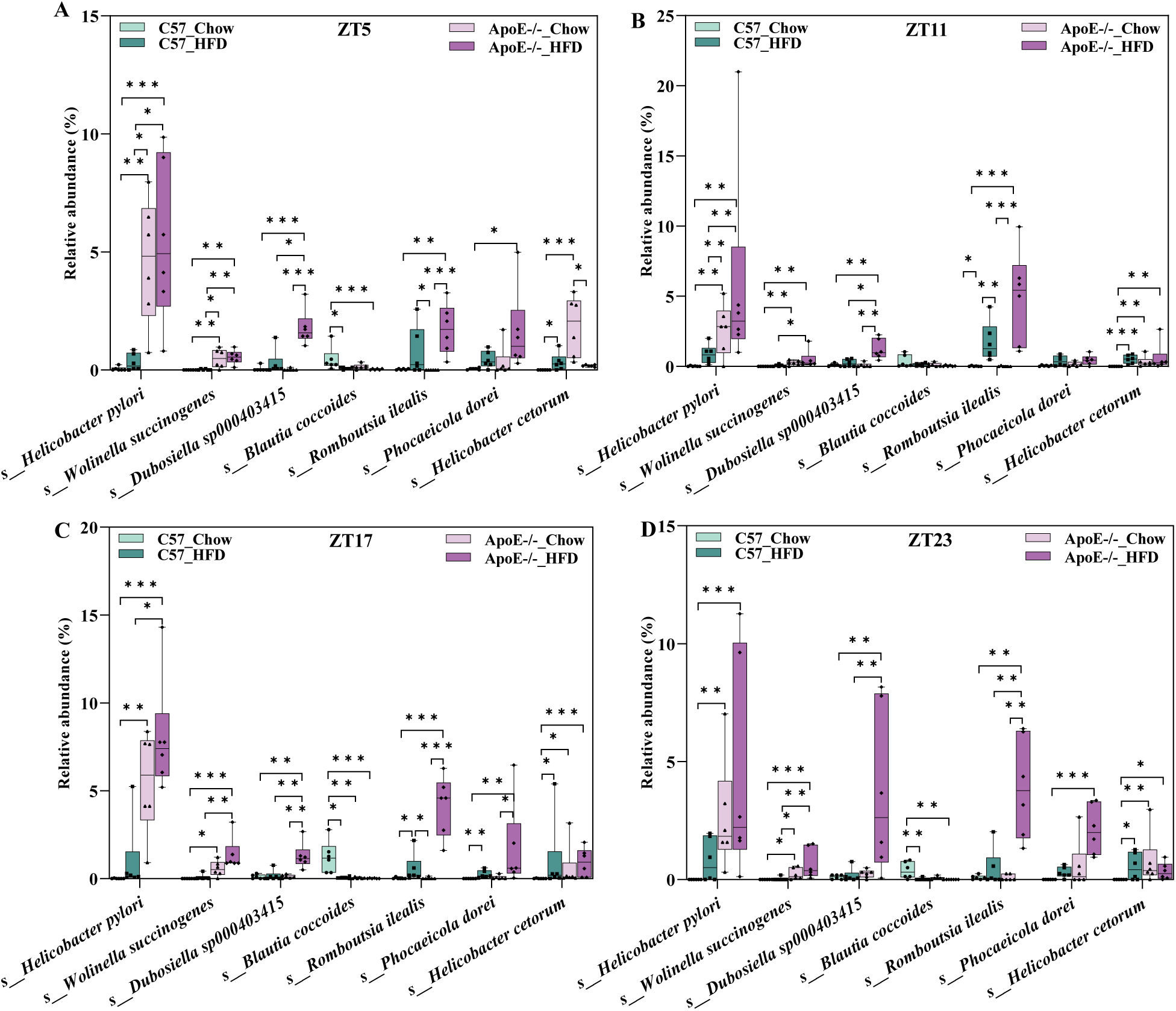
Differences in the relative abundance of species among four groups of mice at ZT5, ZT11, ZT17, and ZT23, respectively. Relative abundance was expressed as mean ± standard error. * indicates *P* < 0.05, ** indicates *P* < 0.01, *** indicates *P* < 0.001; n=6.

**Figure S6.**
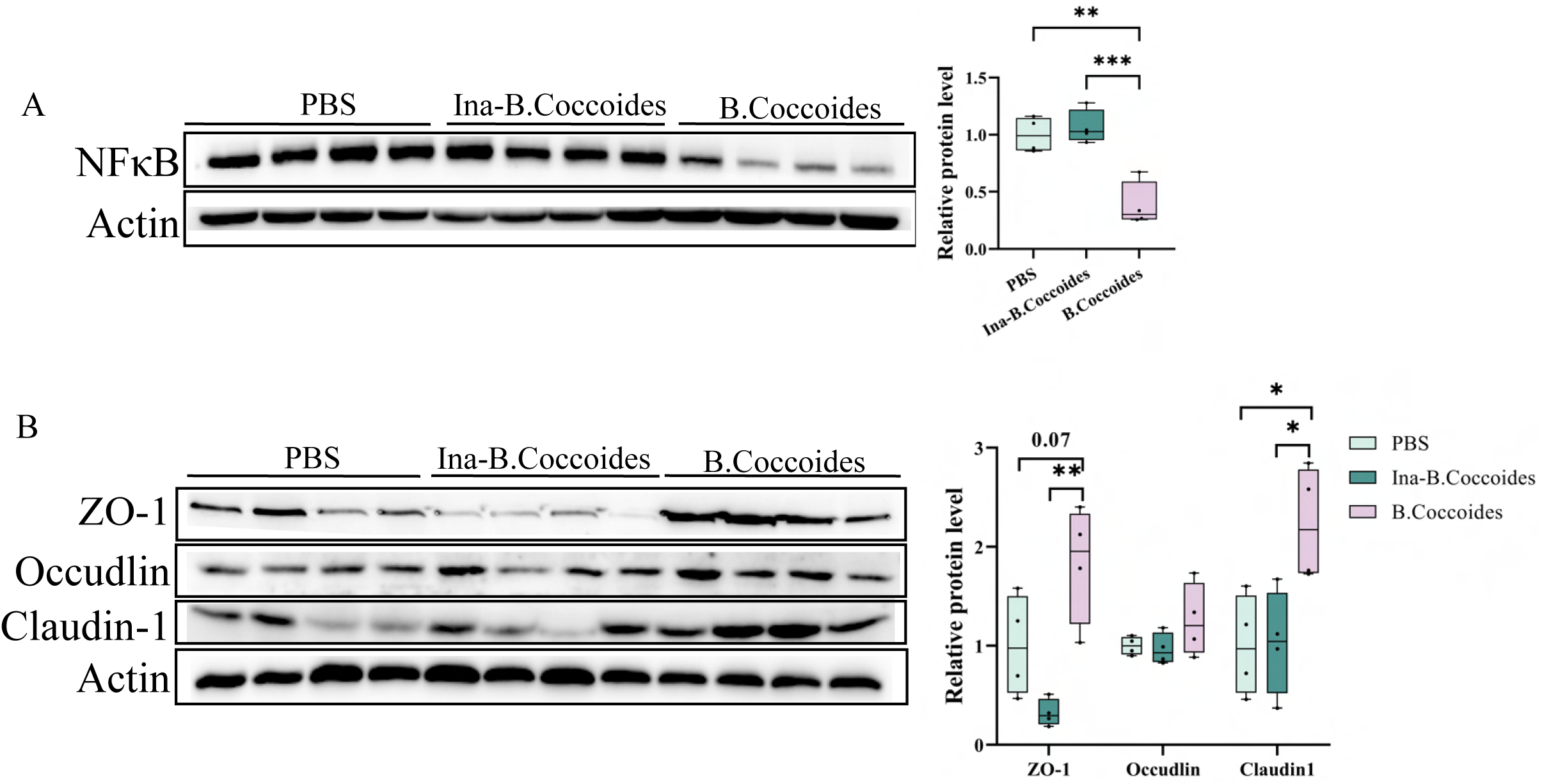
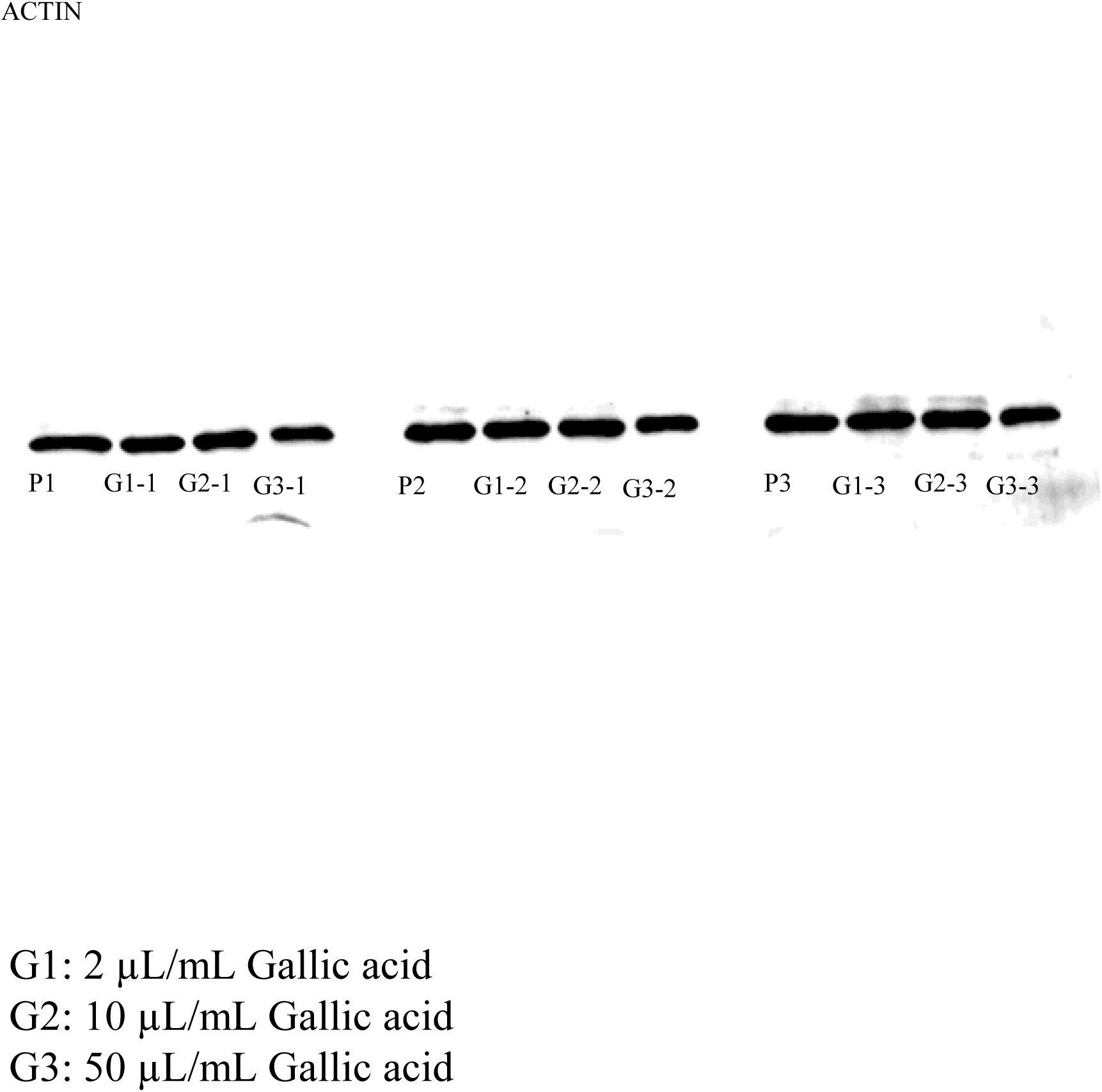
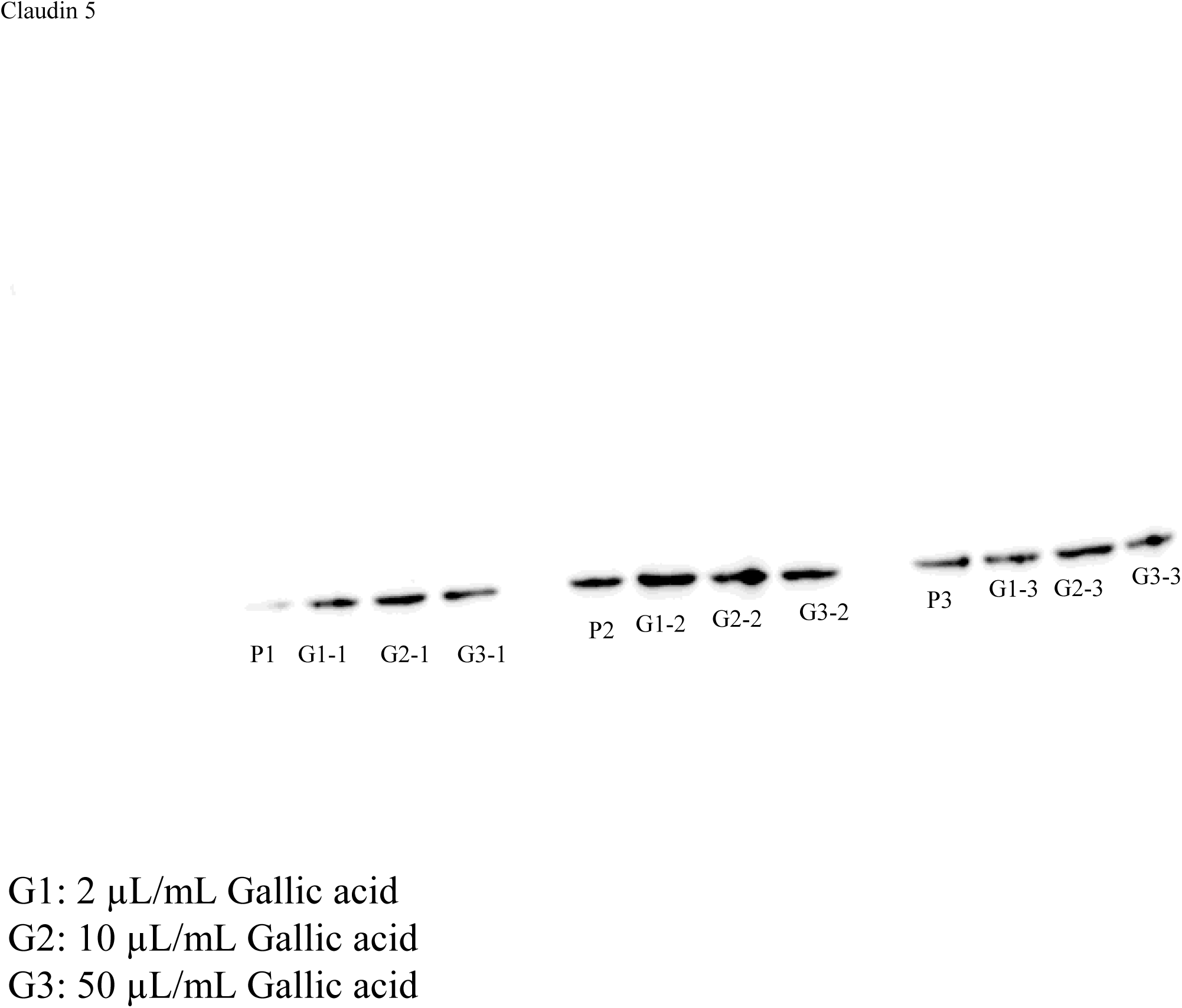
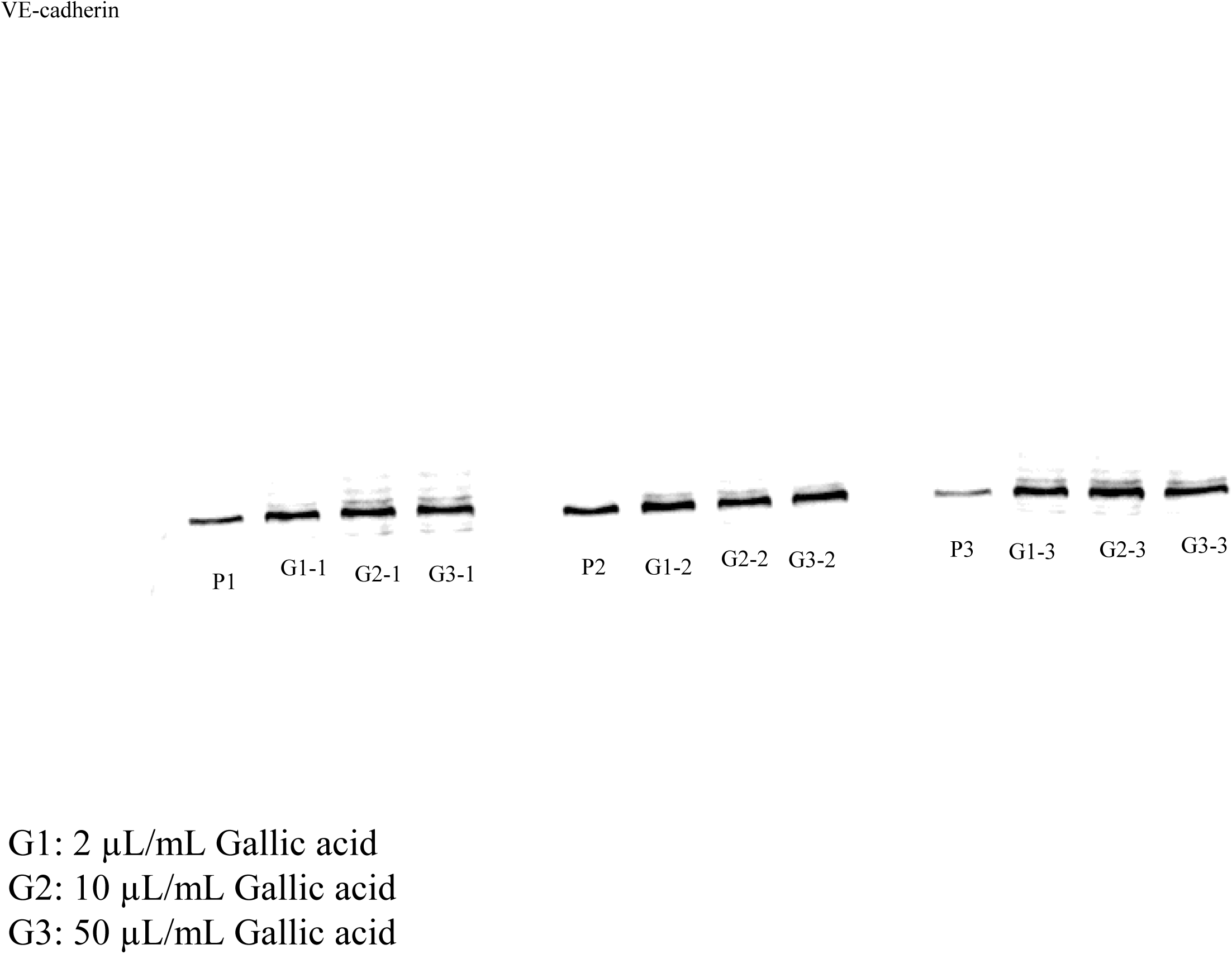
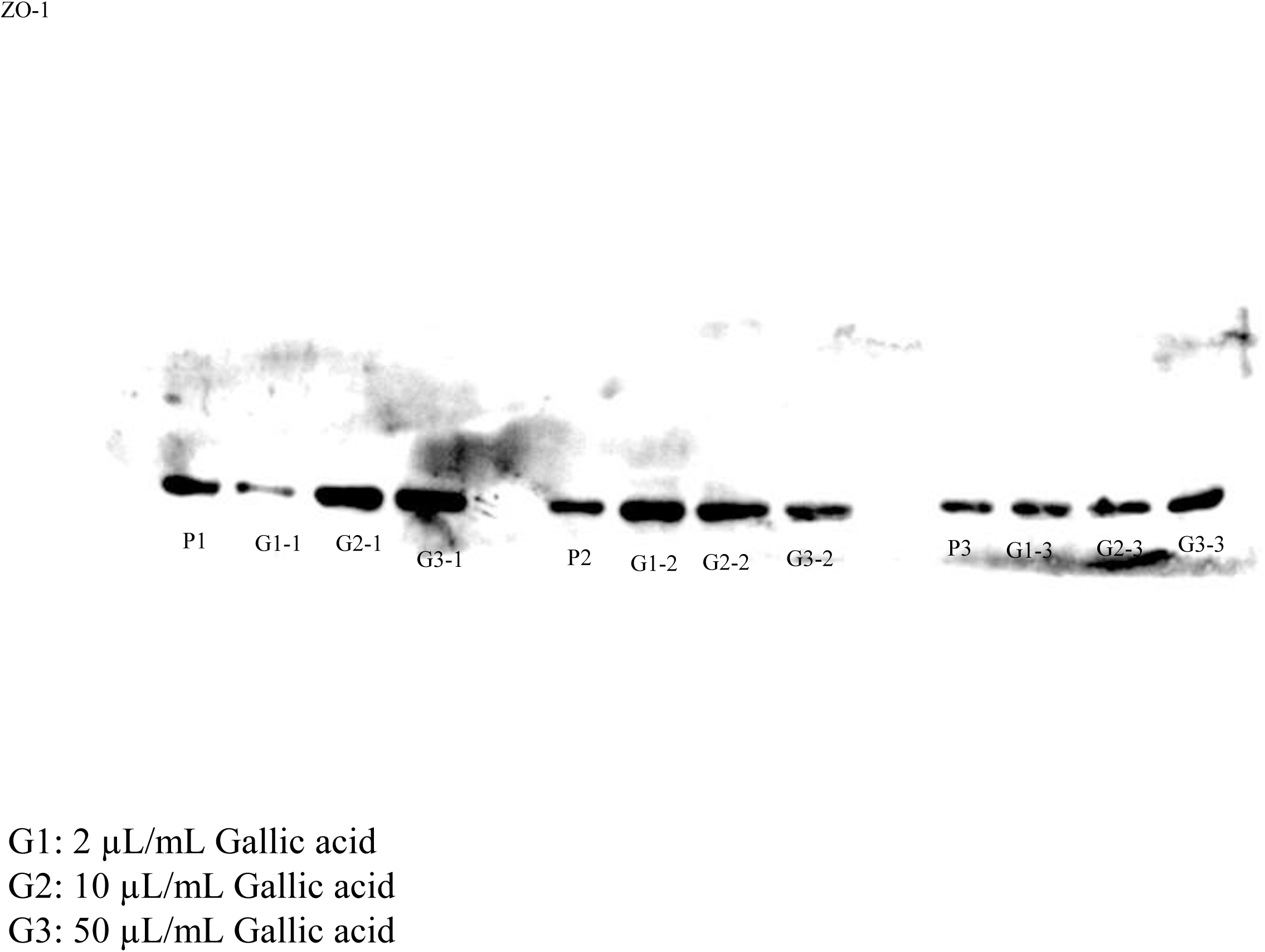
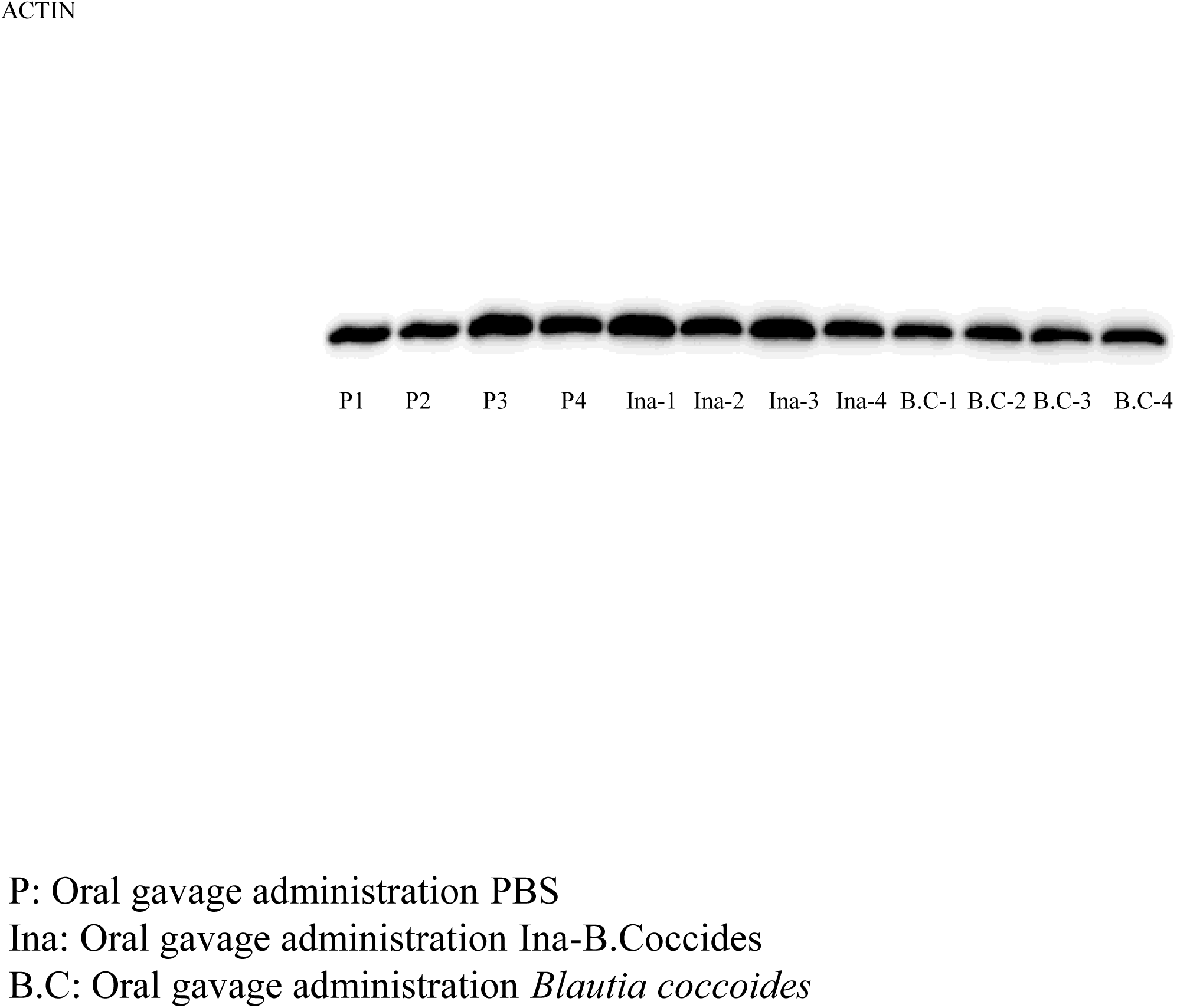
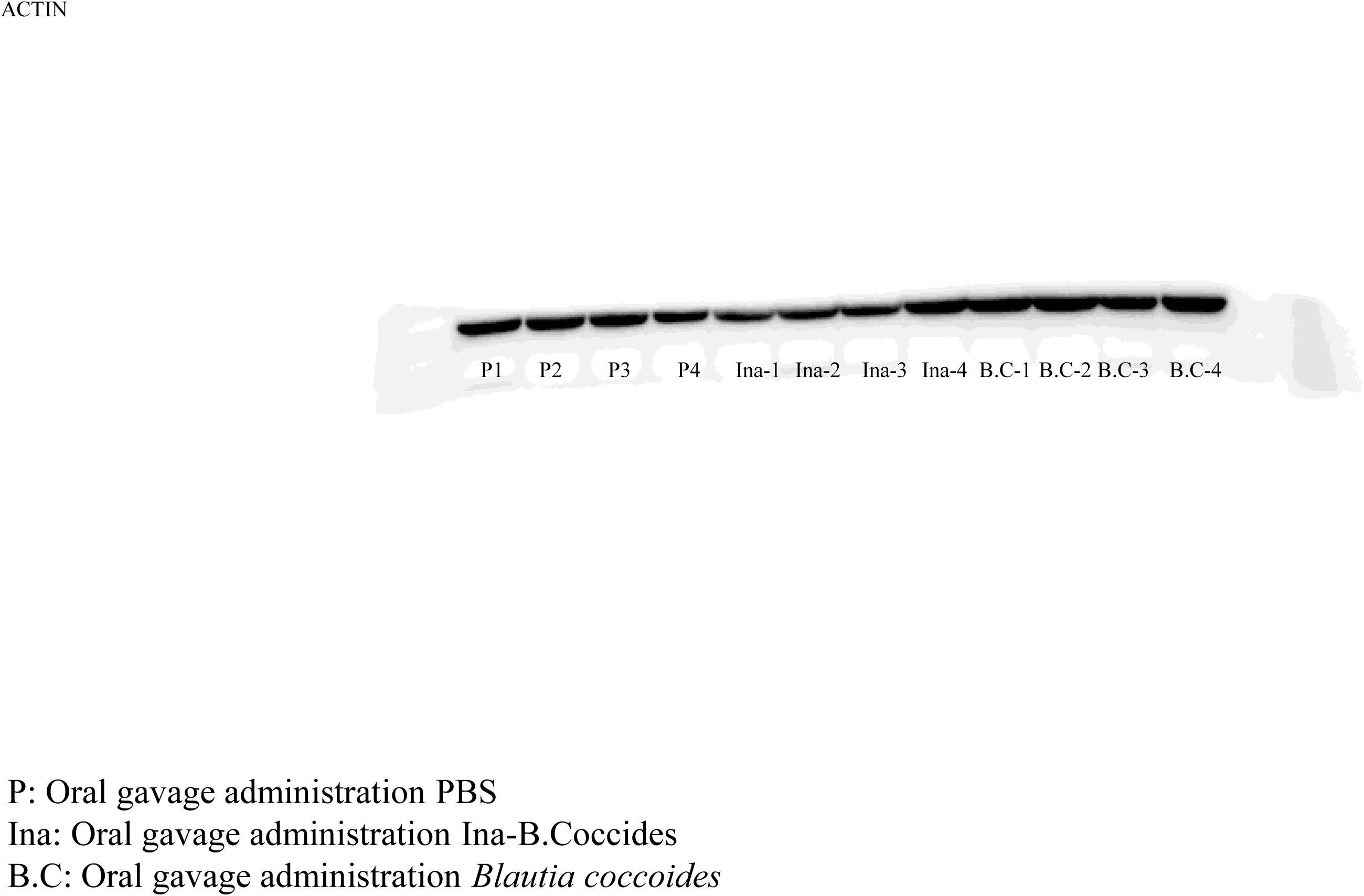
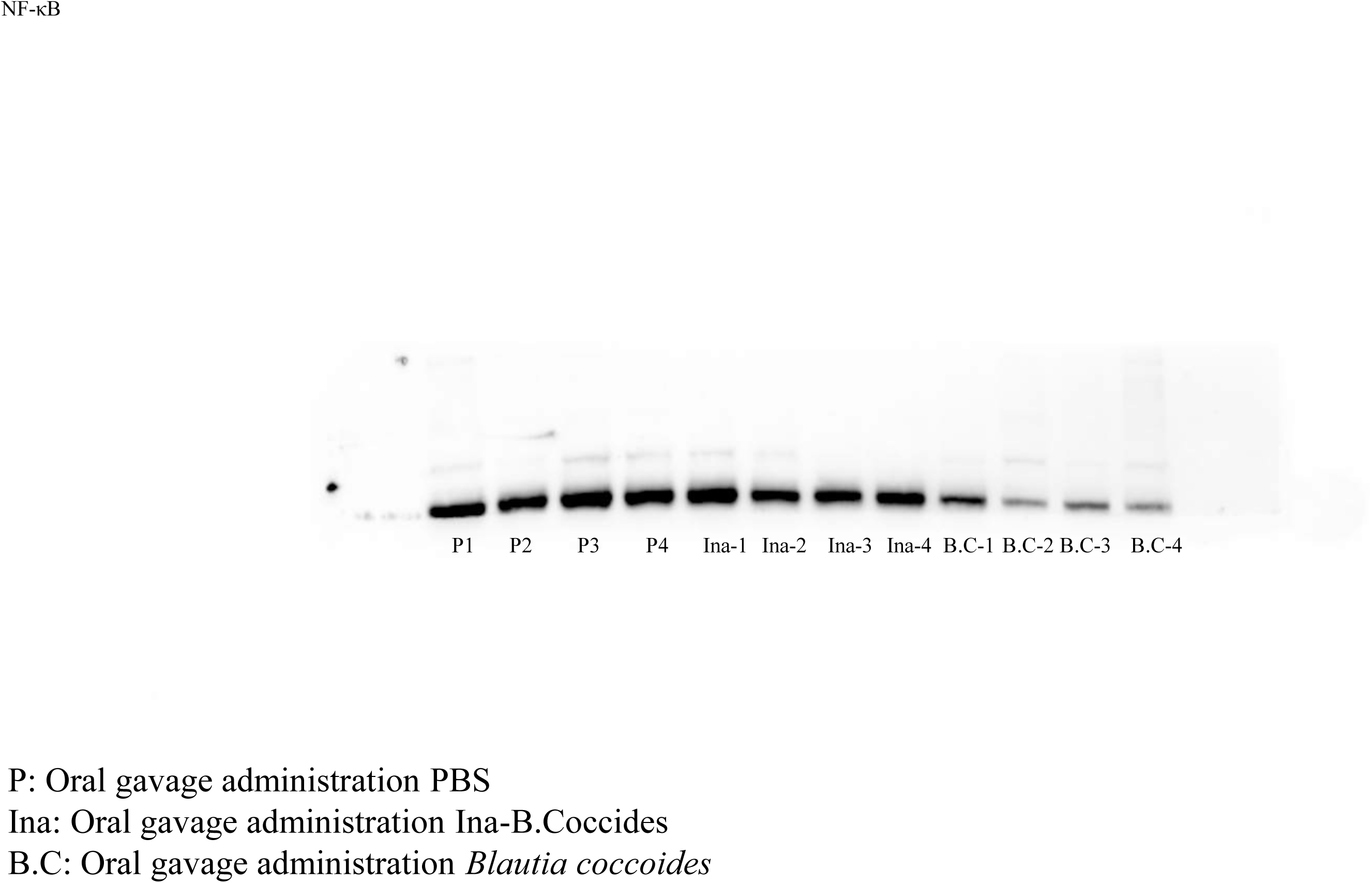
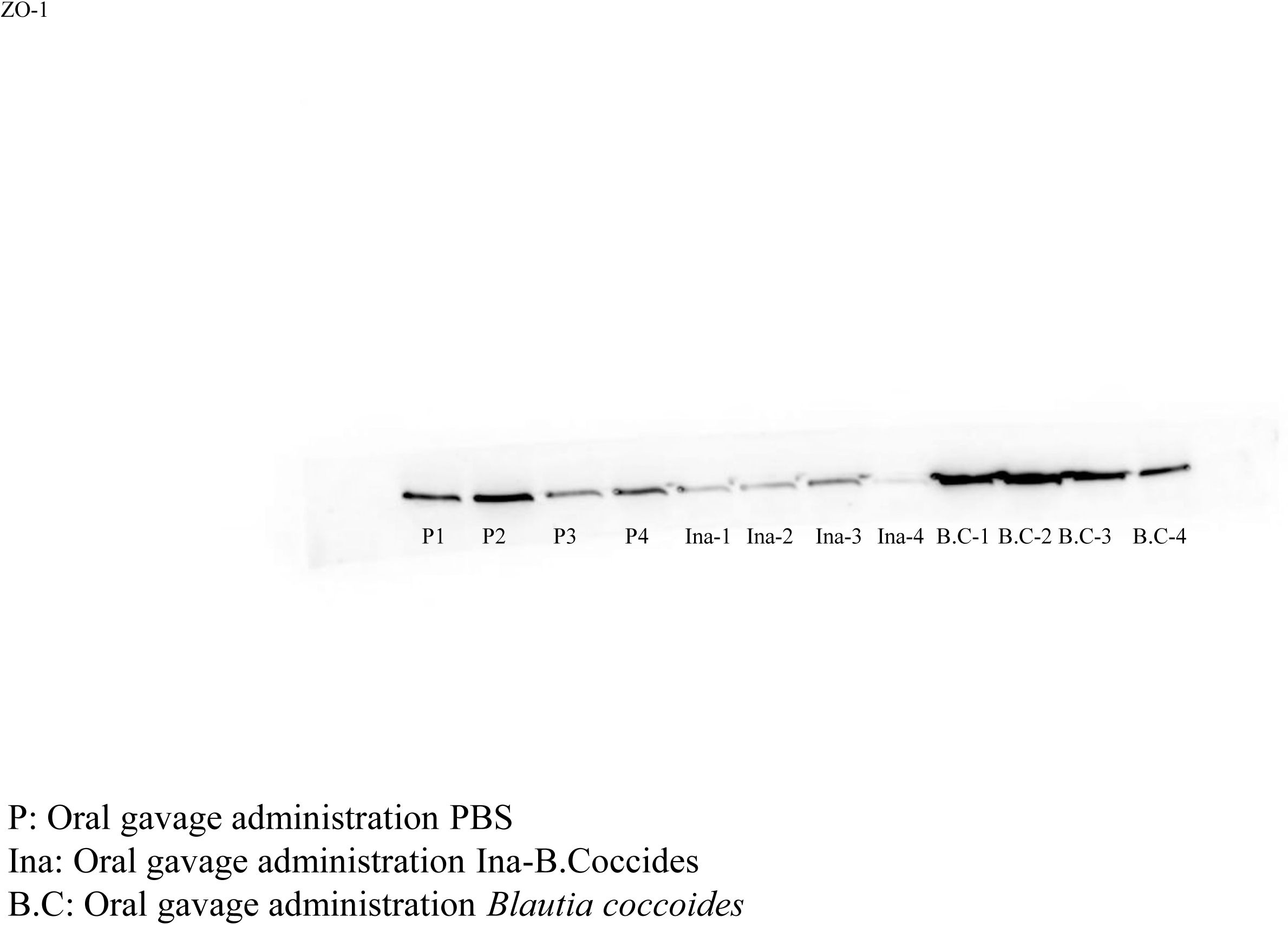
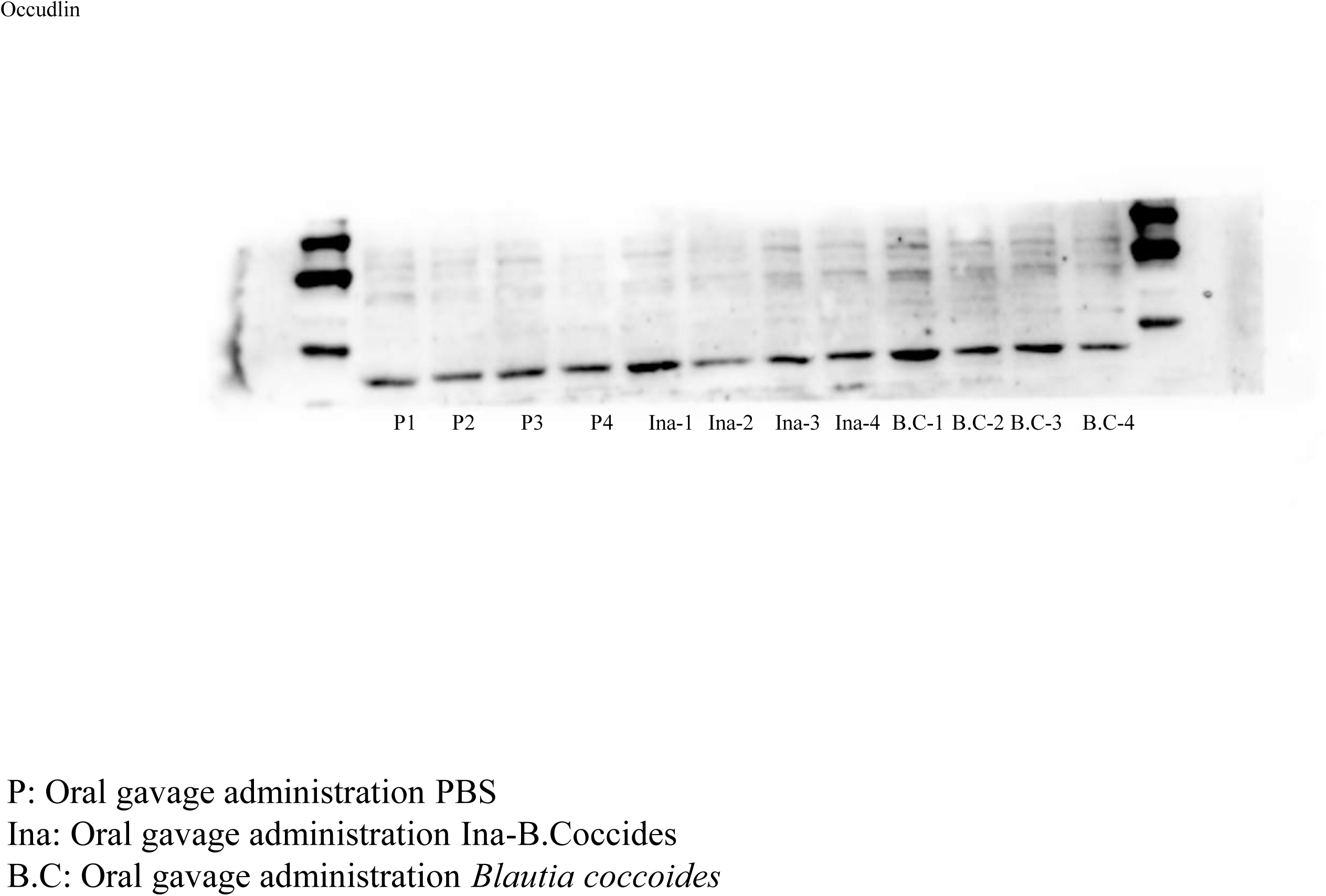
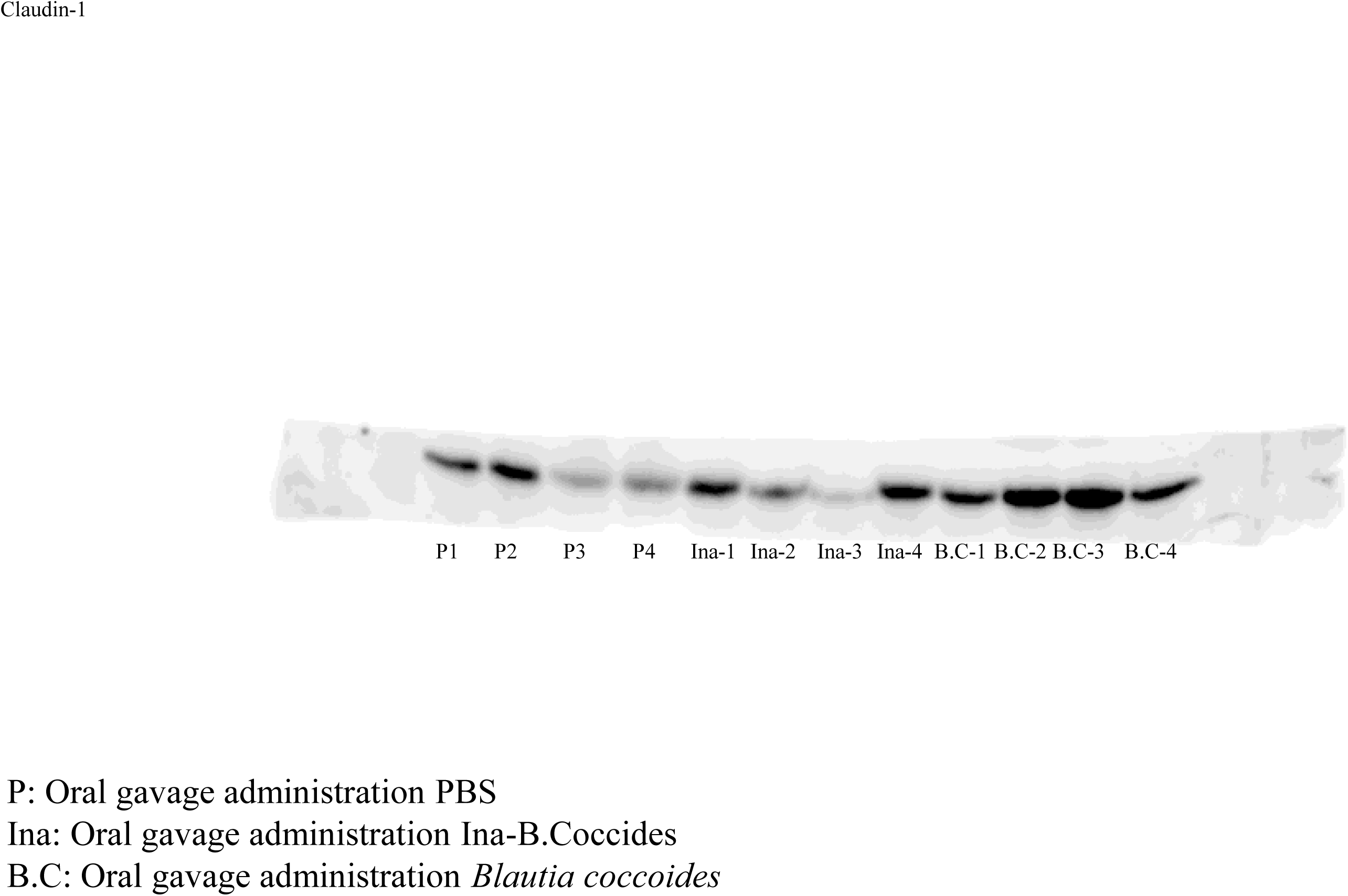
*Blautia coccoides* intervention significantly improved the colon barrier function of ApoE^-/-^ mice with HCHFD. (A) *Blautia coccoides* intervention significantly inhibits the expression level of NF-κB; (B) *Blautia coccoides* intervention significantly increased the expression levels of ZO-1 and Claudin1. Relative abundance was expressed as mean ± standard error. * indicates *P* < 0.05, ** indicates *P* < 0.01, *** indicates *P* < 0.001; n=4.

## Reference

1. Joseph P, Leong D, McKee M, Anand SS, Schwalm JD, Teo K, Mente A, Yusuf S. Reducing the Global Burden of Cardiovascular Disease, Part 1: The Epidemiology and Risk Factors. Circ Res 2017, 121(6): 677-694.

2. Zhao D. Epidemiological Features of Cardiovascular Disease in Asia. **JACC Asia** 2021, 1(1): 1–13.

3. Bjorkegren JLM, Lusis AJ. Atherosclerosis: Recent developments. Cell 2022, 185(10): 1630–1645.

4. Roth GA, Mensah GA, Johnson CO, Addolorato G, Ammirati E, Baddour LM, Barengo NC, Beaton AZ, Benjamin EJ, Benziger CP, Bonny A, Brauer M, Brodmann M, Cahill TJ, Carapetis J, Catapano AL, Chugh SS, Cooper LT, Coresh J, Criqui M, DeCleene N, Eagle KA, Emmons-Bell S, Feigin VL, Fernandez-Sola J, Fowkes G, Gakidou E, Grundy SM, He FJ, Howard G, Hu F, Inker L, Karthikeyan G, Kassebaum N, Koroshetz W, Lavie C, Lloyd-Jones D, Lu HS, Mirijello A, Temesgen AM, Mokdad A, Moran AE, Muntner P, Narula J, Neal B, Ntsekhe M, Moraes de Oliveira G, Otto C, Owolabi M, Pratt M, Rajagopalan S, Reitsma M, Ribeiro ALP, Rigotti N, Rodgers A, Sable C, Shakil S, Sliwa-Hahnle K, Stark B, Sundstrom J, Timpel P, Tleyjeh IM, Valgimigli M, Vos T, Whelton PK, Yacoub M, Zuhlke L, Murray C, Fuster V, Group G-N-JGBoCDW. Global Burden of Cardiovascular Diseases and Risk Factors, 1990-2019: Update From the GBD 2019 Study. J Am Coll Cardiol 2020, 76(25): 2982-3021.

5. Vaduganathan M, Mensah GA, Turco JV, Fuster V, Roth GA. The Global Burden of Cardiovascular Diseases and Risk: A Compass for Future Health. **J Am Coll Cardiol** 2022, 80(25): 2361–2371.

6. Cai YY, Huang FQ, Lao X, Lu Y, Gao X, Alolga RN, Yin K, Zhou X, Wang Y, Liu B, Shang J, Qi LW, Li J. Integrated metagenomics identifies a crucial role for trimethylamine-producing Lachnoclostridium in promoting atherosclerosis. **NPJ Biofilms Microbiomes** 2022, 8(1): 11.

7. Bartolomaeus H, Balogh A, Yakoub M, Homann S, Marko L, Hoges S, Tsvetkov D, Krannich A, Wundersitz S, Avery EG, Haase N, Kraker K, Hering L, Maase M, Kusche-Vihrog K, Grandoch M, Fielitz J, Kempa S, Gollasch M, Zhumadilov Z, Kozhakhmetov S, Kushugulova A, Eckardt KU, Dechend R, Rump LC, Forslund SK, Muller DN, Stegbauer J, Wilck N. Short-Chain Fatty Acid Propionate Protects From Hypertensive Cardiovascular Damage. **Circulation** 2019, 139(11): 1407–1421.

8. Rodriguez-Morato J, Matthan NR. Nutrition and Gastrointestinal Microbiota, Microbial-Derived Secondary Bile Acids, and Cardiovascular Disease. **Curr Atheroscler Rep** 2020, 22(9): 47.

9. Xue HL, Chen X, Yu C, Deng YQ, Zhang Y, Chen S, Chen XC, Chen K, Yang Y, Ling WH. Gut Microbially Produced Indole-3-Propionic Acid Inhibits Atherosclerosis by Promoting Reverse Cholesterol Transport and Its Deficiency Is Causally Related to Atherosclerotic Cardiovascular Disease. **Circulation Research** 2022, 131(5): 404–420.

10. Zhu YJ, Dwidar M, Nemet I, Buffa JA, Sangwan N, Li XMS, Anderson JT, Romano KA, Fu XM, Funabashi M, Wang ZN, Keranahalli P, Battle S, Tittle AN, Hajjar AM, Gogonea V, Fischbach MA, DiDonato JA, Hazen SL. Two distinct gut microbial pathways contribute to meta-organismal production of phenylacetylglutamine with links to cardiovascular disease. **Cell Host & Microbe** 2023, 31(1): 18–32.e9.

11. Bishehsari F, Voigt RM, Keshavarzian A. Circadian rhythms and the gut microbiota: from the metabolic syndrome to cancer. **Nat Rev Endocrinol** 2020, 16(12): 731–739.

12. Choi H, Rao MC, Chang EB. Gut microbiota as a transducer of dietary cues to regulate host circadian rhythms and metabolism. **Nature Reviews Gastroenterology & Hepatology** 2021, 18(10): 679–689.

13. Rothschild D, Weissbrod O, Barkan E, Kurilshikov A, Korem T, Zeevi D, Costea PI, Godneva A, Kalka IN, Bar N, Shilo S, Lador D, Vila AV, Zmora N, Pevsner-Fischer M, Israeli D, Kosower N, Malka G, Wolf BC, Avnit-Sagi T, Lotan-Pompan M, Weinberger A, Halpern Z, Carmi S, Fu J, Wijmenga C, Zhernakova A, Elinav E, Segal E. Environment dominates over host genetics in shaping human gut microbiota. **Nature** 2018, 555(7695): 210-215.

14. David LA, Maurice CF, Carmody RN, Gootenberg DB, Button JE, Wolfe BE, Ling AV, Devlin AS, Varma Y, Fischbach MA, Biddinger SB, Dutton RJ, Turnbaugh PJ. Diet rapidly and reproducibly alters the human gut microbiome. **Nature** 2014, 505(7484): 559-563.

15. Amato KR, Yeoman CJ, Kent A, Righini N, Carbonero F, Estrada A, Gaskins HR, Stumpf RM, Yildirim S, Torralba M, Gillis M, Wilson BA, Nelson KE, White BA, Leigh SR. Habitat degradation impacts black howler monkey (Alouatta pigra) gastrointestinal microbiomes. **ISME J** 2013, 7(7): 1344–1353.

16. Lozupone C, Lladser ME, Knights D, Stombaugh J, Knight R. UniFrac: an effective distance metric for microbial community comparison. **ISME J** 2011, 5(2): 169–172.

17. Ding P, Tong Y, Wu S, Yin X, Liu H, He X, Song Z, Zhang H. The Sexual Effect of Chicken Embryos on the Yolk Metabolites and Liver Lipid Metabolism. **Animals (Basel**) 2021, 12(1):71.

18. Su Y, Luo YH, Zhang LL, Smidt H, Zhu WY. Responses in gut microbiota and fat metabolism to a halogenated methane analogue in Sprague Dawley rats. **Microb Biotechnol** 2015, 8(3): 519–526.

19. Thaiss CA, Zeevi D, Levy M, Zilberman-Schapira G, Suez J, Tengeler AC, Abramson L, Katz MN, Korem T, Zmora N, Kuperman Y, Biton I, Gilad S, Harmelin A, Shapiro H, Halpern Z, Segal E, Elinav E. Transkingdom control of microbiota diurnal oscillations promotes metabolic homeostasis. **Cell** 2014, 159(3): 514–529.

20. Hughes ME, Hogenesch JB, Kornacker K. JTK_CYCLE: an efficient nonparametric algorithm for detecting rhythmic components in genome-scale data sets. **J Biol Rhythms** 2010, 25(5): 372–380.

21. Zhu Y, Dwidar M, Nemet I, Buffa JA, Sangwan N, Li XS, Anderson JT, Romano KA, Fu X, Funabashi M, Wang Z, Keranahalli P, Battle S, Tittle AN, Hajjar AM, Gogonea V, Fischbach MA, DiDonato JA, Hazen SL. Two distinct gut microbial pathways contribute to meta-organismal production of phenylacetylglutamine with links to cardiovascular disease. **Cell Host Microbe** 2023, 31(1): 18–32 e19.

22. Bartolomaeus H, Balogh A, Yakoub M, Homann S, Markó L, Höges S, Tsvetkov D, Krannich A, Wundersitz S, Avery EG, Haase N, Kräker K, Hering L, Maase M, Kusche-Vihrog K, Grandoch M, Fielitz J, Kempa S, Gollasch M, Zhumadilov Z, Kozhakhmetov S, Kushugulova A, Eckardt KU, Dechend R, Rump LC, Forslund SK, Müller DN, Stegbauer J, Wilck N. Short-Chain Fatty Acid Propionate Protects From Hypertensive Cardiovascular Damage. **Circulation** 2019, 139(11): 1407–1421.

23. Wang Z, Klipfell E, Bennett BJ, Koeth R, Levison BS, Dugar B, Feldstein AE, Britt EB, Fu X, Chung YM, Wu Y, Schauer P, Smith JD, Allayee H, Tang WH, DiDonato JA, Lusis AJ, Hazen SL. Gut flora metabolism of phosphatidylcholine promotes cardiovascular disease. **Nature** 2011, 472(7341): 57-63.

24. Wu X, Chen L, Zeb F, Li C, Jiang P, Chen A, Xu C, Haq IU, Feng Q. Clock-Bmal1 mediates MMP9 induction in acrolein-promoted atherosclerosis associated with gut microbiota regulation. **Environ Pollut** 2019, 252(Pt B): 1455–1463.

25. Morris CJ, Purvis TE, Hu K, Scheer FA. Circadian misalignment increases cardiovascular disease risk factors in humans. **Proc Natl Acad Sci U S A** 2016, 113(10): E1402–1411.

26. Schober A, Blay RM, Saboor Maleki S, Zahedi F, Winklmaier AE, Kakar MY, Baatsch IM, Zhu M, Geissler C, Fusco AE, Eberlein A, Li N, Megens RTA, Banafsche R, Kumbrink J, Weber C, Nazari-Jahantigh M. MicroRNA-21 Controls Circadian Regulation of Apoptosis in Atherosclerotic Lesions. **Circulation** 2021, 144(13): 1059–1073.

27. Crnko S, Du Pre BC, Sluijter JPG, Van Laake LW. Circadian rhythms and the molecular clock in cardiovascular biology and disease. **Nat Rev Cardiol** 2019, 16(7): 437–447.

28. Chaix A, Manoogian ENC, Melkani GC, Panda S. Time-Restricted Eating to Prevent and Manage Chronic Metabolic Diseases. **Annu Rev Nutr** 2019, 39: 291–315.

29. Wang P, Song M, Eliassen AH, Wang M, Fung TT, Clinton SK, Rimm EB, Hu FB, Willett WC, Tabung FK, Giovannucci EL. Optimal dietary patterns for prevention of chronic disease. **Nat Med** 2023, 29(3): 719–728.

30. Collaborators GBDD. Health effects of dietary risks in 195 countries, 1990-2017: a systematic analysis for the Global Burden of Disease Study 2017. Lancet 2019, 393(10184): 1958-1972.

31. Dantas Machado AC, Brown SD, Lingaraju A, Sivaganesh V, Martino C, Chaix A, Zhao P, Pinto AFM, Chang MW, Richter RA, Saghatelian A, Saltiel AR, Knight R, Panda S, Zarrinpar A. Diet and feeding pattern modulate diurnal dynamics of the ileal microbiome and transcriptome. **Cell Rep** 2022, 40(1): 111008.

32. Pushpass RAG, Alzoufairi S, Jackson KG, Lovegrove JA. Circulating bile acids as a link between the gut microbiota and cardiovascular health: impact of prebiotics, probiotics and polyphenol-rich foods. **Nutrition Research Reviews** 2022, 35(2): 161–180.

33. Cortes V, Eckel RH. Insulin and Bile Acids in Cholesterol Homeostasis: New Players in Diabetes-Associated Atherosclerosis. **Circulation** 2022, 145(13): 983–986.

34. Nishijima S, Suda W, Oshima K, Kim SW, Hirose Y, Morita H, Hattori M. The gut microbiome of healthy Japanese and its microbial and functional uniqueness. **DNA Res** 2016, 23(2): 125–133.

35. Niu YG, Hu XM, Song YL, Wang CC, Luo PX, Ni SH, Jiao FX, Qiu J, Jiang WH, Yang S, Chen J, Huang R, Jiang HZ, Chen SH, Zhai QW, Xiao J, Guo FF. Blautia Coccoides is a Newly Identified Bacterium Increased by Leucine Deprivation and has a Novel Function in Improving Metabolic Disorders. **Advanced Science** 2024, 11(18):e2309255.

36. Liu X, Mao B, Gu J, Wu J, Cui S, Wang G, Zhao J, Zhang H, Chen W. Blautia-a new functional genus with potential probiotic properties? **Gut Microbes** 2021, 13(1): 1–21.

37. Holmberg SM, Feeney RH, Vishnu PPK, Puértolas-Balint F, Singh DK, Wongkuna S, Zandbergen L, Hauner H, Brandl B, Nieminen AIJNC. The gut commensalBlautiamaintains colonic mucus function under low-fiber consumption through secretion of short-chain fatty acids. **Nat Commun** 2024, 15(1):3502.

38. Deaver JA, Eum SY, Toborek M. Circadian Disruption Changes Gut Microbiome Taxa and Functional Gene Composition. **Front Microbiol** 2018, 9: 737.

39. Nishino K, Nishida A, Inoue R, Kawada Y, Ohno M, Sakai S, Inatomi O, Bamba S, Sugimoto M, Kawahara M, Naito Y, Andoh A. Analysis of endoscopic brush samples identified mucosa-associated dysbiosis in inflammatory bowel disease. **J Gastroenterol** 2018, 53(1): 95–106.

40. Meier KHU, Julian T, Hai L, Melanie L, Tobias F, Nicola Z, Shinichi S, Macpherson AJ, Uwe SJNM. Metabolic landscape of the male mouse gut identifies different niches determined by microbial activities. **Nat Metab** 2023, 5(6): 968–980.

41. Patel SS, Goyal RK. Cardioprotective effects of gallic acid in diabetes-induced myocardial dysfunction in rats. **Pharmacognosy Res** 2011, 3(4): 239–245.

42. Bai J, Zhang Y, Tang C, Hou Y, Ai X, Chen X, Zhang Y, Wang X, Meng X. Gallic acid: Pharmacological activities and molecular mechanisms involved in inflammation-related diseases. **Biomed Pharmacother** 2021, 133: 110985.

43. Jadon A, Bhadauria M, Shukla S. Protective effect of Terminalia belerica Roxb. and gallic acid against carbon tetrachloride induced damage in albino rats. **J Ethnopharmacol** 2007, 109(2): 214–218.

44. Wu Y, Li K, Zeng M, Qiao B, Zhou B. Serum Metabolomics Analysis of the Anti-Inflammatory Effects of Gallic Acid on Rats With Acute Inflammation. **Front Pharmacol** 2022, 13: 830439.

45. Iwona C, Katharina UJIJoIC, Research M. TNF-α in the cardiovascular system: from physiology to therapy. Int. J. Interferon, Cytokine Mediator Res 2015, 7: 9–25.

46. Adjuto-Saccone M, Soubeyran P, Garcia J, Audebert S, Tournaire RJCD, Disease. TNF-α induces endothelial–mesenchymal transition promoting stromal development of pancreatic adenocarcinoma. **Cell Death Dis** 2021, 12(7):649.

47. Otani T, Furuse M. Tight Junction Structure and Function Revisited: (Trends in Cell Biology 30, 805-817, 2020). Trends Cell Biol 2020, 30(12): 1014.

48. Cong X, Kong W. Endothelial tight junctions and their regulatory signaling pathways in vascular homeostasis and disease. **Cell Signal** 2020, 66:109485.

